# The Human Mitochondrial Genome Encodes for an Interferon-Responsive Host Defense Peptide

**DOI:** 10.1101/2023.03.02.530691

**Authors:** MC Rice, M Imun, SW Jung, CY Park, JS Kim, RW Lai, CR Barr, JM Son, K Tor, E Kim, RJ Lu, I Cohen, BA Benayoun, C Lee

**Author notes:** Correspondence (C.L.), (B.A.B). Department of Food & Nutrition, College of Health Science, The University of Suwon, Hwaseong-si, 18323, Korea (CY Park). These authors contributed equally (MC Rice, M Imun, SW Jung).

## Abstract

The mitochondrial DNA (mtDNA) can trigger immune responses and directly entrap pathogens, but it is not known to encode for active immune factors. The immune system is traditionally thought to be exclusively nuclear-encoded. Here, we report the identification of a mitochondrial-encoded host defense peptide (HDP) that presumably derives from the primordial proto-mitochondrial bacteria. We demonstrate that MOTS-c (mitochondrial open reading frame from the twelve S rRNA type-c) is a mitochondrial-encoded amphipathic and cationic peptide with direct antibacterial and immunomodulatory functions, consistent with the peptide chemistry and functions of known HDPs. MOTS-c targeted *E. coli* and methicillin-resistant *S. aureus* (MRSA), in part, by targeting their membranes using its hydrophobic and cationic domains. In monocytes, IFNγ, LPS, and differentiation signals each induced the expression of endogenous MOTS-c. Notably, MOTS-c translocated to the nucleus to regulate gene expression during monocyte differentiation and programmed them into macrophages with unique transcriptomic signatures related to antigen presentation and IFN signaling. MOTS-c-programmed macrophages exhibited enhanced bacterial clearance and shifted metabolism. Our findings support MOTS-c as a first-in-class mitochondrial-encoded HDP and indicates that our immune system is not only encoded by the nuclear genome, but also by the co-evolved mitochondrial genome.

## Introduction

The endosymbiotic theory posits that mitochondria originate from once free-living bacteria that infected eukaryotic ancestral cells. Indeed, owing to their bacterial origin^1^, mitochondria still retain several prokaryotic characteristics, including N-formylated proteins and a circular DNA, that register as quasi-self damage-associated molecular patterns (DAMPs)^2^. Multiple pattern-recognition receptors (PRRs) can sense DAMPs and elicit pro-inflammatory and type I interferon responses^2^. Further, mitochondrial DNA (mtDNA) can be actively released by neutrophils^3^ and eosinophils^4^ to physically target pathogens and by lymphocytes to signal type I interferon responses^5^.

While mtDNA itself can trigger immune responses and directly entrap pathogens, it is not known to encode for genes that yield immune factors. The continuous expansion of our proteome, powered by the characterization of short/small open reading frames (sORFs) in the nuclear^6,7^ and mitochondrial^8,9^ genomes that yield functional peptides, provides unprecedented opportunities for the discovery of host defense peptides (HDPs), also known as antimicrobial peptides (AMPs)^10,11^. All kingdoms of life possess a vast repertoire of AMPs that are now pursued therapeutically as antibiotics, antivirals, cell-penetrating peptides, immunomodulators, and cancer targeting peptides^10,12–18^. HDPs are microproteins of 10 to 50 amino acids in length^10,12,14^. We have previously identified a mitochondrial-encoded sORF, MOTS-c (mitochondrial ORF from the twelve S rRNA type-c), which yields a 16 amino acid peptide^19^. MOTS-c has a key role in regulating cellular homeostasis under cellular stress and during aging^20^, in part, by directly translocating to the nucleus to regulate adaptive gene expression^19,21–24^.

Based on the endosymbiosis theory, early communication between the proto-mitochondria and proto-eukaryotic cell likely occurred on an immunological basis with immune factors that were encoded within both genomes. Today, bacteria still use gene-encoded peptides, known as bacteriocins, that regulate vital cellular processes (*e.g.* ribosomal processes, DNA replication, and cell wall synthesis^25^) to control inter- and intra-species proliferation^16,26–29^. In the unique endosymbiotic context, these peptides may have served to not only protect the newly formed union from other pathogens, but also to regulate the growth and metabolism of the opposing quasi-self (*i.e.* the present-day mitochondria and nucleus). Thus, mitochondrial-derived peptides (MDPs)^30^, including MOTS-c, may inherently possess immuno-metabolic functions and represent a primordial arm of the eukaryotic immune system (**Figure 1A**) that evolved with dual roles as cellular regulators during aging^19,20,31^.

**Figure 1.**
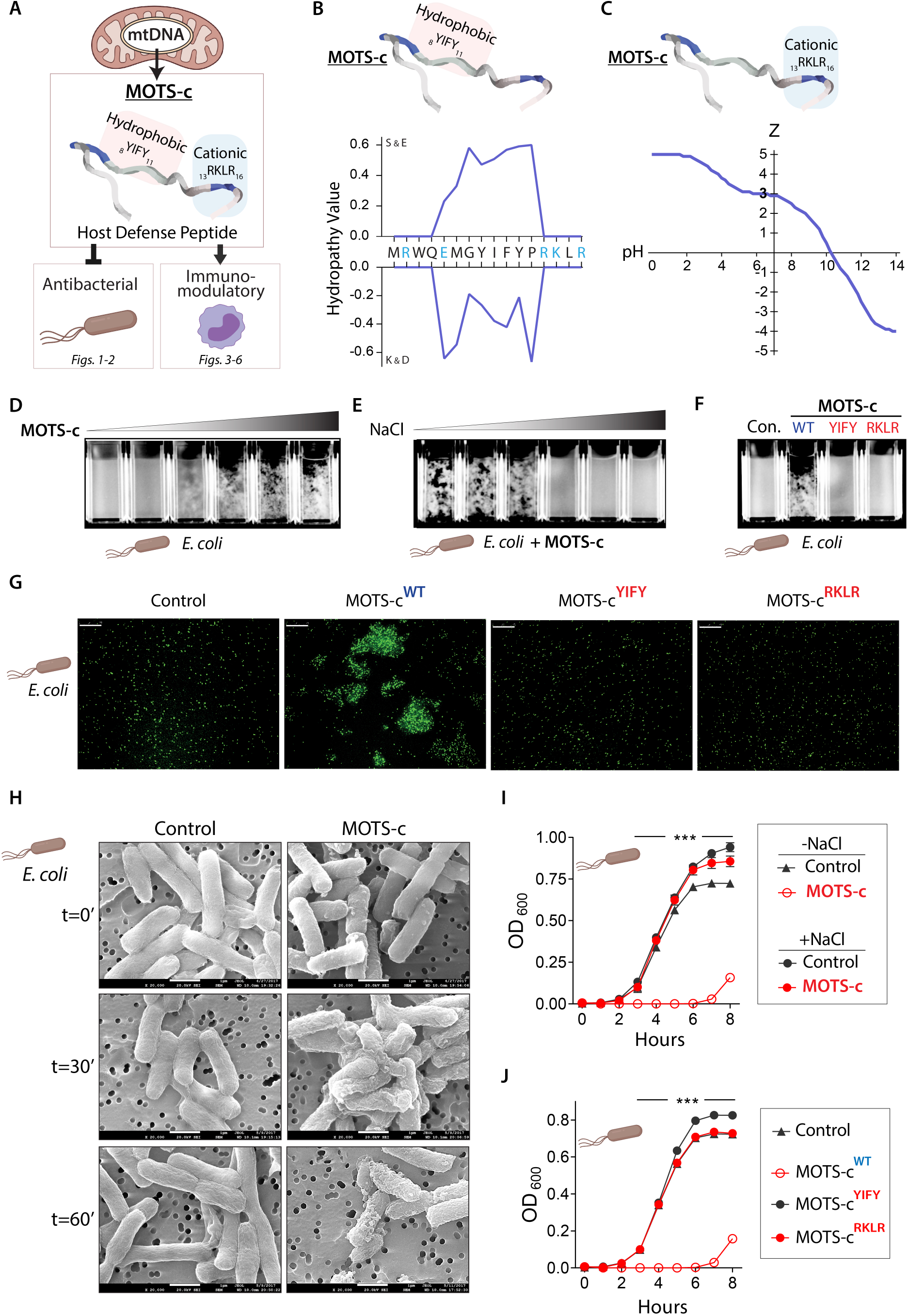
MOTS-c is a mitochondrial-encoded host defense peptide (HDP). (**A**) Bacteria and bacteria-derived mitochondria possess gene-encoded immune peptides, known as host defense peptides (HDPs) in higher eukaryotes. (**B**) MOTS-c has a hydrophobic core (_8_YIFY_11_), determined using the hydrophobicity scales of Kyte and Doolittle (K&D)^32^ and Sweet and Eisenberg (S&E)^33^. Blue: cationic residues. (**C**) MOTS-c has a cationic tail (_13_RKLR_16_) that confers positive charge (Z) across a pH range. (**D**) MOTS-c treatment (0-100 µM) immediately aggregates *E. coli* in a dose dependent manner. (**E**) MOTS-c-dependent *E. coli* aggregation is lost in increasing salt concentrations (NaCl; 0-1%) and (**F, G**) requires its hydrophobic core and cationic tail, consistent with other HDPs. EGFP-expressing *E. coli* (BL21) shown. Wildtype MOTS-c (WT) and mutants devoid of its hydrophobic (YIFY: _8_YIFY_11_>_8_AAAA_11_) or cationic domain (RKLR: _13_RKLR_16_>_13_AAAA_16_). Bar, 75 µm. (**H**) Scanning electron micrographs of *E. coli* treated with MOTS-c (100 µM) for 0 (immediate fixation), 30, and 60 minutes (n=3). Representative images shown. Bar, 100 nm. (**I-J**) Growth curve of *E. coli* (BL21), measured by optical density at 600 nm (OD_600_), following (n=6) (**I**) MOTS-c treatment in the presence of 1% NaCl, and (**J**) treatment with wildtype (WT) MOTS-c and mutants devoid of its hydrophobic (_8_YIFY_11_>_8_AAAA_11_; YIFY), or cationic domain (_13_RKLR_16_>_13_AAAA_16_; RKLR). Data expressed as mean +/- SEM. Two-way ANOVA repeated measures. *** P<0.001.

Here, we report, for the first time, that the human mitochondrial genome encodes for a HDP that directly targets bacteria and regulates monocyte function. This indicates that the human immune system is encoded in both of our co-evolved mitochondrial and nuclear genomes.

## Results

### The mitochondrial-derived peptide MOTS-c compromises bacterial viability in vitro

HDPs are characterized by their cationic and amphipathic residues, typically with a net positive charge. Peptide analysis using the hydrophobicity scales of Kyte and Doolittle^32^ and Sweet and Eisenberg^33^ indicated that MOTS-c is an amphipathic peptide consisting of a core that is enriched with hydrophobic residues (_8_YIFY_11_) (**Figure 1B**). MOTS-c retains a +3 charge under physiological pH, largely owing to the basic tail residues (_13_RKLR_16_) (**Figure 1C**), which may also serve as a heparin-binding domain (XBBXB-like moiety; B: basic residues) that can bind to cell wall carbohydrates (*e.g.* glucan, mannan) and mediate bacterial recognition and binding^34,35^.

Direct bacterial targeting is a hallmark of HDPs^18,36^. Indeed, MOTS-c dynamically associated with bacteria immediately upon contact (**Figures 1D**-**1H and S1**). We mixed MOTS-c (100 µM) with *E. coli* suspended in water. MOTS-c levels in the media (water) decreased concomitantly with increased detection in cell lysates, indicating direct interaction with bacteria (**Figure S1A**). HDPs can cause bacterial aggregation, which immobilizes them to the local infection site and enhances pathogen clearance^37–41^. Consistently, MOTS-c immediately aggregated *E. coli* and MRSA (methicillin-resistant *S. aureus*) in a dose- and growth phase-dependent manner (**Figures 1D and S1B-1C**; **Video S1**). HDPs engage with bacteria through ionic and hydrophobic interactions owing to their hydrophobic and charged residues^42,43^. MOTS-c was ineffective in aggregating bacteria under higher salt concentrations (0-1% in ddH_2_O)(**Figure 1E**), which disrupts ionic interactions and HDP activity^43,44^. The loss of the hydrophobic and cationic domains of MOTS-c, by substituting the residues with alanine (*i.e.* _8_YIFY_11_>_8_AAAA_11_ and _13_RKLR_16_>_13_AAAA_16_), prevented bacterial aggregation (**Figures 1F**-**1G and S1D**), consistent with the importance of these residues for HDP function^10,45^. MOTS-c did not aggregate *S. typhimurium* or *P. aeruginosa* (**Figure S2**), indicating target selectivity independent of Gram status.

HDPs can perturb bacterial membranes via several mechanisms, including progressive blebbing, budding, and pore formation^46–48^. Using scanning electron microscopy (SEM), we visualized a time-dependent progression of MOTS-c-dependent membrane blebbing in *E. coli* (**Figure 1H**) and MRSA (**Figure S3**). Membrane destabilization by HDPs can also cause bacterial aggregation^49–51^. We confirmed rapid compromise of bacterial membrane integrity upon MOTS-c treatment using a fluorescent nucleic acid stain that only penetrates permeabilized membranes^52^ (**Figure S4A**). Further, MOTS-c treatment depleted cellular ATP levels in *E. coli* (**Figure S4B**). Consistently, real-time metabolic flux analyses revealed that MOTS-c treatment perturbed respiration and glycolysis (**Figure S4C**), which requires intact membrane function^53^.

We next confirmed the antimicrobial effect of MOTS-c on bacterial growth. A single treatment of MOTS-c significantly retarded *E. coli* proliferation in liquid culture in a dose-dependent manner (**Figure S4D-S4E**). Two intermediate doses of MOTS-c (50 µM) had comparable effects to a single high-dose treatment (100 µM) (**Figure S4D**), reflecting the significance of antibiotic treatment frequency in addition to absolute dose. Notably, human HDPs can reach high intracellular concentrations compared to their low circulating levels: LL-37 can reach 40 µM^54–57^ and defensins can constitute 5-7% of the total protein content of neutrophils^58,59^. Further, HDPs can reach very high membrane-bound concentrations that are 10,000 times of aqueous solutions (80 mM)^36^. Interruption of ionic interaction, achieved by higher salt concentration (**Figure 1I**) or loss of the hydrophobic or cationic domains by alanine-substitution mutagenesis (*i.e.* _8_YIFY_11_>_8_AAAA_11_ and _13_RKLR_16_>_13_AAAA_16_; 100 µM) (**Figure 1J**), fully reversed the antimicrobial function of MOTS-c on *E. coli* growth. Multiple HDPs also have intracellular roles that contribute to the antimicrobial effect^60–63^. Using an inducible MOTS-c expression vector, we found that the endogenous overexpression of MOTS-c significantly inhibited *E. coli* (BL21) growth (**Figure S4F**), indicating a two-stage mechanism of targeting membranes and intracellular components.

### MOTS-c enhances survival from MRSA exposure in vivo

To confirm the sustained antibacterial effects of MOTS-c *in vivo*, we inoculated female mice with MRSA that had been treated with or without MOTS-c (**Figure 2A-2O**). While intraperitoneal inoculation of MRSA was lethal, MOTS-c-treated MRSA was not (16.67% *vs.* 100% survival, respectively) (**Figure 2A**). To further explore whether lethal peritonitis could be induced by high levels of pathogen-associated molecular patterns (PAMPs) alone or if live bacteria were necessary, we simultaneously inoculated a third group of mice with heat-killed MRSA. All mice in this group survived, indicating that live bacteria are required to cause lethal peritonitis (**Figure 2A**). Notably, both the heat-killed MRSA and MOTS-c-treated MRSA groups showed complete survival; however, the group that received MOTS-c-treated MRSA lost weight during the 72-hour infection, suggesting a distinct response to the inoculation (**Figure 2B**). MOTS-c-treated MRSA, sampled from the inoculated preparation (6×10^8^ CFU), showed a 3.4-fold reduction in CFU on LB agar plates (**Figure 2C**). This suggests that treatment with MOTS-c reduced the MRSA bacterial burden sufficiently for the mice to clear the infection and recover. To determine the level of inflammatory response to MRSA inoculation, we measured circulating cytokines (IL-1β, IL-2, IL-4, IL-5, IL-6, IL-10, IL-12p70, IFN-γ, and TNF-α) 6 hours post-infection. Indeed, mice inoculated with MOTS-c-treated MRSA had an intermediary induction of cytokines between those of the control and heat-killed MRSA groups (**Figure 2D**-**2L**). At 72 hours post-infection, cytokine levels in the MOTS-c-treated MRSA group were significantly reduced and closer to those of the heat-killed group compared to the control group (sole surviving mouse), indicating resolution of inflammation (**Figure 2D**-**2L**). Elevated blood urea nitrogen (BUN), aspartate transaminase (AST), and alanine transaminase (ALT) levels in circulation indicate kidney and liver injury during lethal infection^64^. After 72 hours, mice inoculated with MOTS-c-treated MRSA had lower BUN levels and no difference in AST and ALT levels compared to the heat-killed MRSA group (**Figure 2M**-**2O**). This suggests that the MOTS-c-treated MRSA group induced an inflammatory immune response without significant organ damage. These results were consistent in male mice, with 20% *vs.* 100% survival in the control and MOTS-c-treated group, respectively, and a 19.8-fold reduction in CFU of sampled inoculated MRSA preparation (4×10^8^ CFU) on LB agar plates (**Figure S5**). This indicates that, similar to other known HDPs, the antimicrobial effects of MOTS-c depend on the stoichiometric ratio of MOTS-c:MRSA and their proximity^65^. These results suggest that MOTS-c can opsonize bacteria, perhaps for immune clearance, and neutralize their pathogenicity.

**Figure 2.**
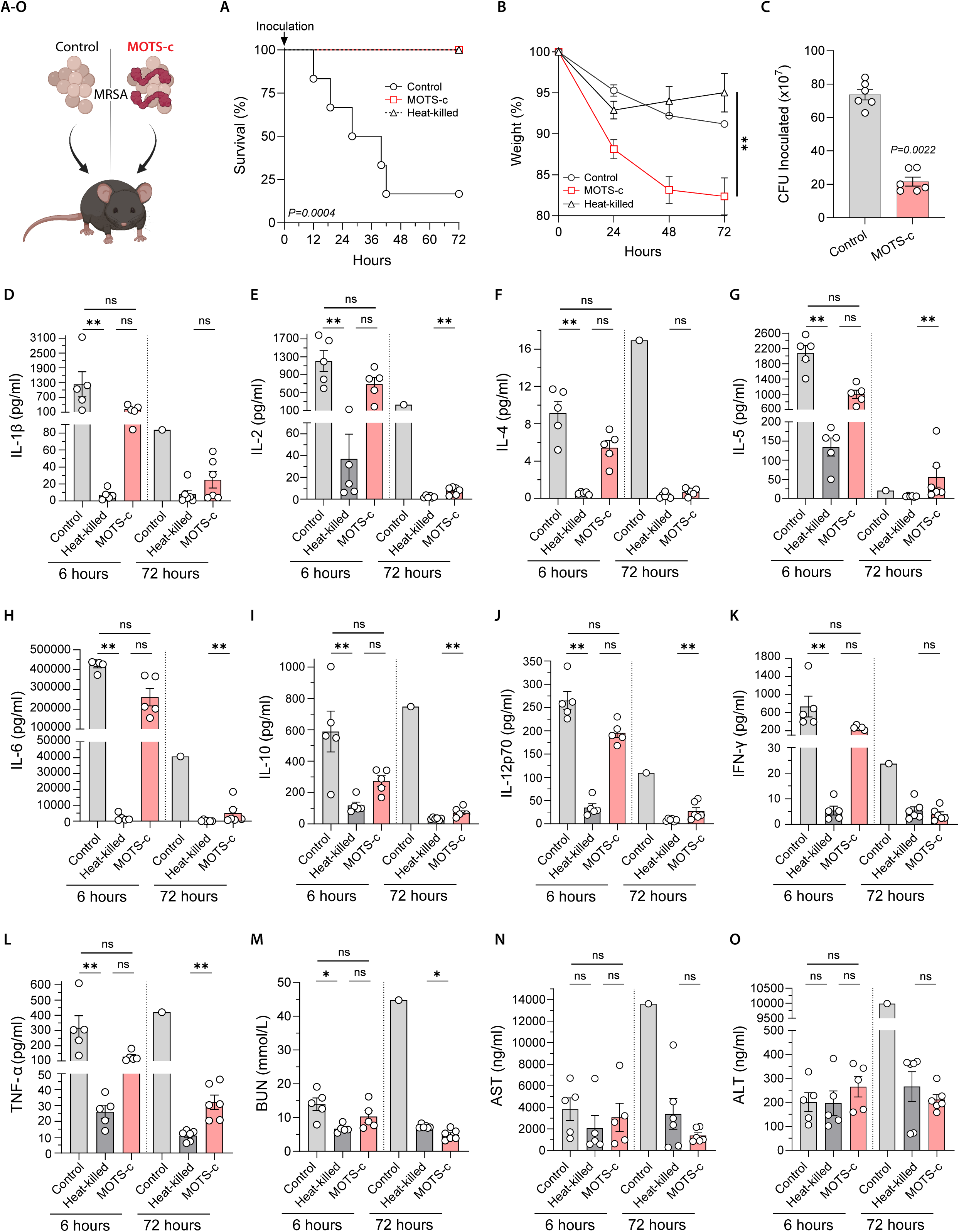
MOTS-c enhances survival from MRSA exposure *in vivo*. (**A-O**) 6×10^8^ CFU of mid-log phase MRSA either resuspended in (i) 100 µM MOTS-c, (ii) vehicle (water) or (iii) in vehicle and then heat-killed in a water bath. MRSA preparations were then immediately injected IP into 6-month-old female C57BL/6J mice (n=5-6). Mice were euthanized and blood collected after 6 and 72 hours. (**A**) Survival and (**B**) weight were monitored for 72 hours. (**C**) 6×10^8^ CFU of MRSA resuspended in 100 µM MOTS-c or water was serial diluted and plated on LB agar before injection and colonies counted after overnight incubation. (**D-L**) Cytokines (IL-1β, IL-2, IL-4, IL-5, IL-6, IL-10, IL-12p70, IFN-γ, and TNF-α) were measured in plasma by multiplex ELISA after 6 hours (n=5) and 72 hours (n=1 for control and n=6 for others; surviving mice). Plasma levels of (**M**) Blood urea nitrogen (BUN), (**N**) aspartate transaminase (AST), and (**O**) alanine transaminase (ALT) were measured by ELISA at 6 hours (n=5) and 72 hours (n=1 for control and n=6 for others). Data expressed as mean +/- SEM. Log-rank (Mantel-Cox) test for (A), 2-way ANOVA (repeated measures) for (B), Mann-Whitney test for (C), and Kruskal-Wallis test (6 hour timepoint) and Mann-Whitney test (72 hour timepoint; due to only 1 control surviving.) for (D-O).* P<0.05, ** P<0.01.

### Endogenous MOTS-c is induced upon human monocyte activation in vitro

Although the initial focus on HDP research was on their direct antimicrobial effect, recent studies have established them as key regulators of immune responses, including monocyte activation and differentiation^10,11^. Because (i) the discovery of MOTS-c was inspired by a prior study demonstrating interferon gamma (IFNγ)-induced transcripts from the mitochondrial rRNA genes in monocytes^66^ and (ii) MOTS-c regulates adaptive cellular responses to various types of stress^19,21–24^, we hypothesized that it may act as a modulator of monocyte activation and differentiation. Endogenous MOTS-c levels dynamically increased in a time-dependent manner upon monocyte-to-macrophage differentiation in (i) primary human peripheral blood monocytes by M-CSF (macrophage colony stimulating factor)(**Figure 3A**) and (ii) a human monocytic cell line (THP-1) by phorbol myristate acetate (PMA)^67^(**Figure 3B**). Since they recapitulated the dynamics of MOTS-c upon differentiation signals and they are a tractable cell line, we decided to perform follow-up experiments primarily in the THP-1 system. Endogenous MOTS-c was also induced in THP-1 monocytes in a time-dependent manner following stimulation with lipopolysaccharides (LPS) and IFNγ (**Figure 3C**), a combination of bacterial-derived and cytokine-dependent signals known to synergistically activate monocytes^68–74^. We then tested whether LPS and IFNγ could each induce MOTS-c expression separately. LPS alone increased MOTS-c expression in THP-1 monocytes (**Figure 3D**) and differentiated THP-1 macrophages (**Figure S6A**), consistent with known HDPs^75–83^. IFNγ alone also induced MOTS-c expression in THP-1 monocytes (**Figure 3E**), consistent with the strong induction of transcripts from the mitochondrial rRNA genes in interferon-induced monocyte-like cells^66^. Cellular levels of induced MOTS-c in THP-1 macrophages appear to reach high concentrations (**Figure S6B**), consistent with known HDPs^54–59^.

**Figure 3.**
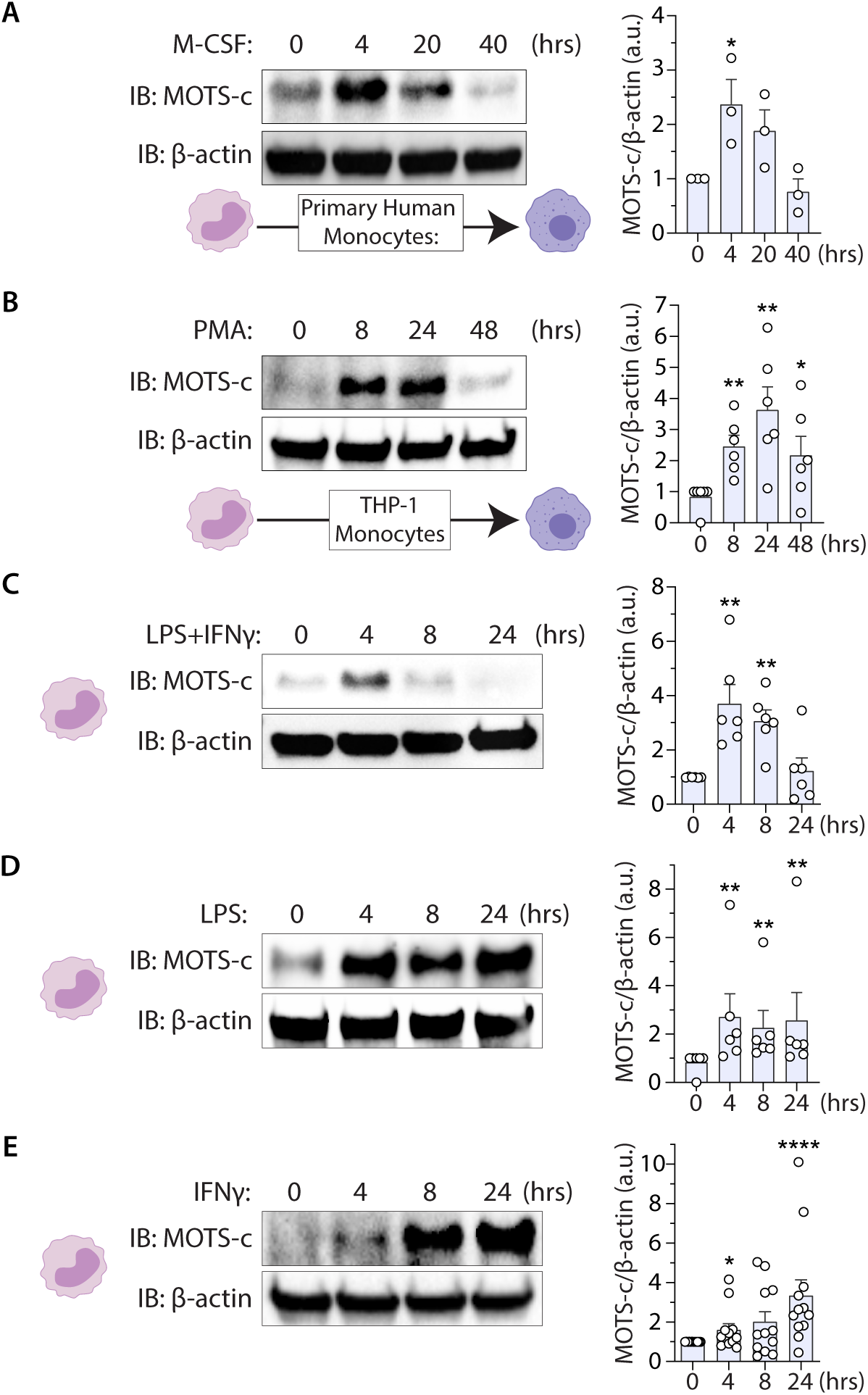
MOTS-c is induced in activated and differentiating monocytes. (**A-B**) Total endogenous MOTS-c levels measured as a function of time following monocyte differentiation in (**A**) primary human monocytes by M-CSF (100 ng/ml) (n=3) and (**B**) THP-1 cells by PMA (15 nM) (n=6). (**C-E**) Total endogenous MOTS-c levels following THP-1 monocyte activation by LPS (100 ng/ml) and IFNγ (20ng/ml) (**C**) in combination (n=6), (**D**) LPS alone (n=6), and (**E**) IFNγ alone (n=12). Data expressed as mean +/- SEM. Mann-Whitney test. *P<0.05, ** P<0.01, **** P<0.0001.

### MOTS-c regulates the differentiation trajectory of human monocytes in vitro

Because HDPs can regulate monocyte activation and differentiation^84–87^, we next tested whether MOTS-c can modulate monocyte differentiation. We previously reported that MOTS-c translocates to the nucleus upon cellular stress to regulate a range of nuclear genes, indicating that mitochondrial-encoded factors can regulate the nuclear genome^21–23,31^. Indeed, endogenous MOTS-c translocated to the nucleus upon differentiation of THP-1 monocytes by PMA in a time-dependent manner (**Figure 4A**) and following LPS+IFNγ stimulation (**Figure S7A**). Consistent with our previous reports^19,21–24^, exogenously treated MOTS-c readily entered THP-1 monocytes (**Figure S7B**); the uptake was significantly retarded at a lower temperature (4°C)(**Figure S7C**), indicating the potential involvement of active transport^88^. Exogenously treated MOTS-c peptide also showed strong nuclear localization (**Figures 4B and S7D**). To test whether MOTS-c regulates nuclear transcriptional programming during early monocyte differentiation, we performed bulk RNA-seq on THP-1 monocytes that were treated with PMA or PMA+MOTS-c for 2 hours (**Figure 4C**-**4H**). Multidimensional scaling (MDS)^89^ revealed that the overall expression profiles of PMA and PMA+MOTS-c monocytes were distinct, as they were clearly separated in the MDS 2-dimensional space (**Figure 4C**). Notably, the separation of the two groups occurred mostly on a single dimension (*i.e.* dimension 1, which captures the largest proportion of variance among samples), compatible with the notion that MOTS-c treatment led to acceleration in differentiation-related gene expression changes. MOTS-c differentially regulated 945 genes between PMA *vs*. PMA+MOTS-c groups (FDR<5%) (**Figure 4D**; **Table S1**), comparable to LL-37 which can affect the expression of >900 nuclear genes in human monocytes^10^.

**Figure 4.**
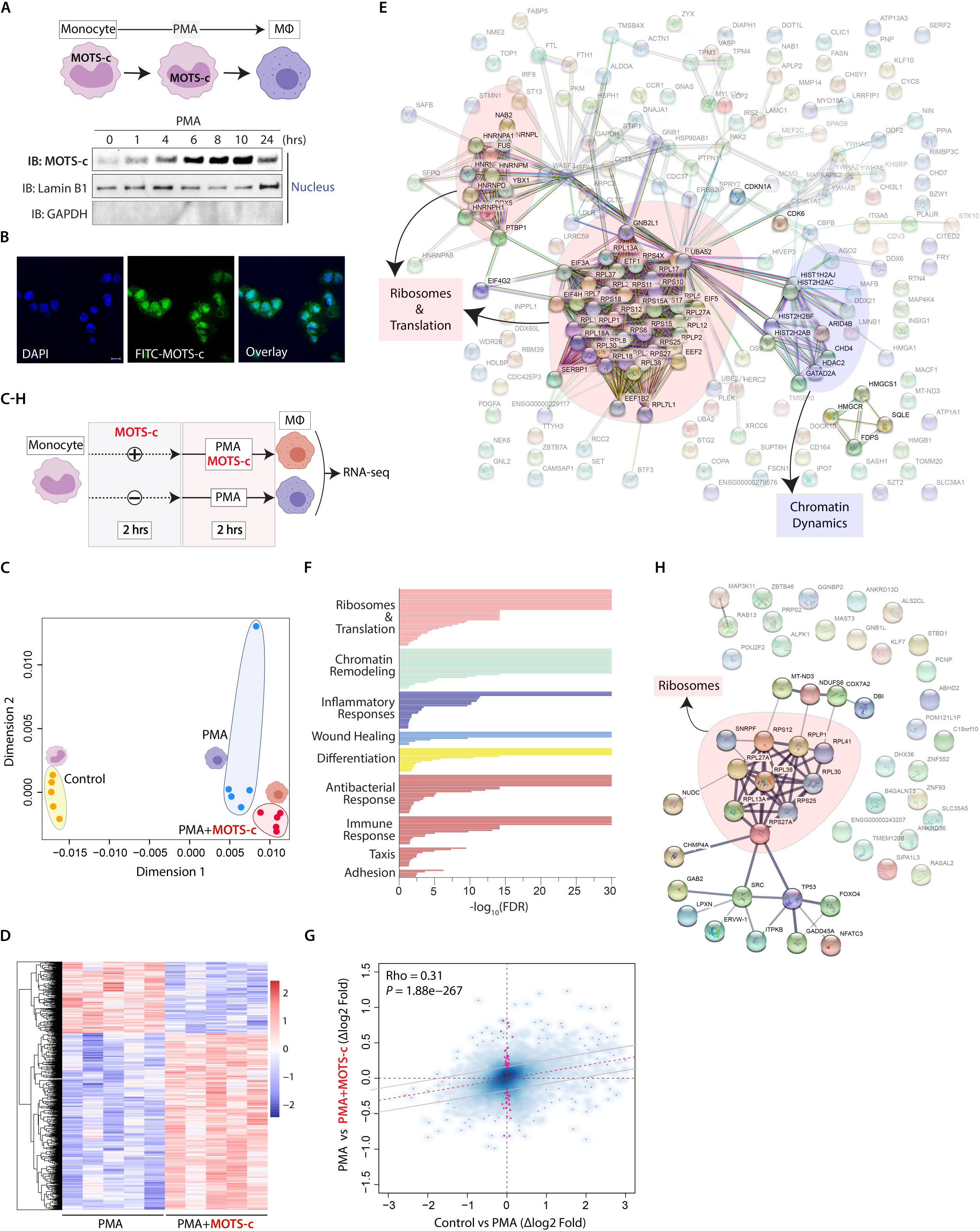
MOTS-c reprograms early nuclear gene expression during monocyte differentiation. (**A**) A time-course measurement of endogenous MOTS-c in purified nuclear extracts following THP-1 monocyte differentiation by PMA (15 nM). (**B**) Confocal fluorescence images of THP-1 monocytes treated with FITC-MOTS-c (1 µM) for 30 minutes, showing nuclear localization. Nucleus marked by DAPI staining. Bar, 10 µm. (**C-H**) THP-1 Monocytes were primed with MOTS-c (10 µM) or vehicle control for 2 hours, then differentiated with PMA +/- MOTS-c (10 µM) for 2 hours, at which time RNA was collected for bulk RNA-seq analysis (n=6); false discovery rate (FDR) < 5%. (**C**) Multidimensional scaling (MDS) analysis across control, PMA, and PMA+MOTS-c groups based on RNA-seq expression profiles after DESeq2 VST normalization. (**D**) Heatmap of significantly differentially regulated genes by MOTS-c by DESeq2 analysis. (**E**) Protein-protein interaction network analysis based on genes that were significantly differentially up- and down-regulated by MOTS-c (FDR < 5%) using the STRING (Search Tool for the Retrieval of Interacting Genes/Proteins) database version 11.0^90^. (**F**) Significantly enriched biological functions based on gene set enrichment analysis (GSEA) using gene ontology (GO). Selected groups are shown, full data in **Table S1**. (**G**) Correlation plot of gene expression changes by DESeq2 upon (i) *regular* induction of differentiation (control *vs.* PMA) compared to (ii) MOTS*-*c-directed induction of differentiation (PMA *vs*. PMA+MOTS-c). Spearman Rank correlation (Rho), and significance of this correlation are reported. Genes that are significantly regulated only upon MOTS-c treatment but not during normal differentiation are highlighted in red and may underlie a specific MOTS-c-induced macrophage state. (**H**) Protein-protein interaction network analysis based on the 64 genes that were significantly differentially up- and down-regulated by MOTS-c (FDR < 5%) as described in (G) using the STRING database version 11.0 ^90^. MΦ=macrophage.

Using the STRING (Search Tool for the Retrieval of Interacting Genes/Proteins) database version 11.0^90^, which can assess putative changes in protein-protein interaction networks based on our RNA-seq analysis, we identified large gene clusters in the PMA+MOTS-c group, compared to the PMA group, that were related to (i) ribosomes and translation initiation and (ii) chromatin dynamics (**Figure 4E**; full results in **Table S2**). Consistently, functional enrichment analysis for Gene Ontology (GO) gene sets revealed that the most significantly targeted functions by MOTS-c included (i) ribosomal and translational processes and (ii) chromatin dynamics (selected terms in **Figure 4F**; full results in **Table S2**). We then compared gene expression changes during normal differentiation (control *vs.* PMA) and MOTS-c-programmed differentiation (PMA *vs*. PMA+MOTS-c) and Spearman rank correlation (Rho) analysis revealed that changes were significantly correlated, consistent with the notion that MOTS-c treatment can accelerate the normal macrophage differentiation program (**Figure 4G**). In this context, we identified 64 genes that were specifically attributed to MOTS-c regulation after PMA induction (**Figure 4G**; **Table S1D**), of which ribosomal genes were again represented based on STRING analysis (**Figure 4H**). Consistently, ribosomal levels and differential translation are known to regulate cellular differentiation and lineage commitment^91–94^, indicating that MOTS-c may broadly impact gene expression during monocyte differentiation.

### In vitro MOTS-c-treated human monocytes yield functionally distinct macrophages

Due to the impact of MOTS-c on the transcriptome of THP-1 monocytes, we then asked whether MOTS-c can regulate monocyte differentiation to produce macrophages that are functionally distinct. A single exposure to MOTS-c during differentiation enhanced monocyte adherence, an important step for extravasation and differentiation to macrophages^95^. We found that a greater number of MOTS-c-programmed primary human monocytes became adherent 3 days (**Figure 5A**) and 6 days (**Figure S8A**) after M-CSF stimulation, indicating an accelerated differentiation program, consistent with our RNA-seq analysis (**Figure 4C and 4G**). Next, we exposed THP-1 monocytes to MOTS-c at the onset of differentiation and functionally characterized the MOTS-c-programmed macrophages 4 days later. First, MOTS-c-programmed THP-1 macrophages exhibited a significant increase in bacterial killing capacity, determined using a gentamicin protection assay (**Figure 5B**). Second, we characterized the differential response of MOTS-c-programmed macrophages to LPS stimulation, including cytokine secretion/expression and cellular metabolism (**Figure 5C**-**5F**). MOTS-c-programmed THP-1 macrophages exhibited (i) a shift in LPS-induced secretion of IL-1β, IL-1Ra and TNFα as measured by ELISA (**Figure 5C**) and (ii) selective impact on LPS-induced chemokine and cytokine gene expression as measured by RT-qPCR (**Figures 5D and S8B**), and (iii) suppressed oxygen consumption in response to LPS, a metabolic reflection of macrophage activity^96–100^, following acute stimulation (**Figure 5E**) and later reduced spare respiratory capacity 16 hours post-stimulation (**Figure 5F**). These results are in line with the previously described impact of LL-37, a well-described human HDP, in reprogramming monocytes to differentiate into macrophages with enhanced antibacterial capacity^101^, suggesting that the mitochondrial-encoded MOTS-c may act in a similar fashion.

**Figure 5.**
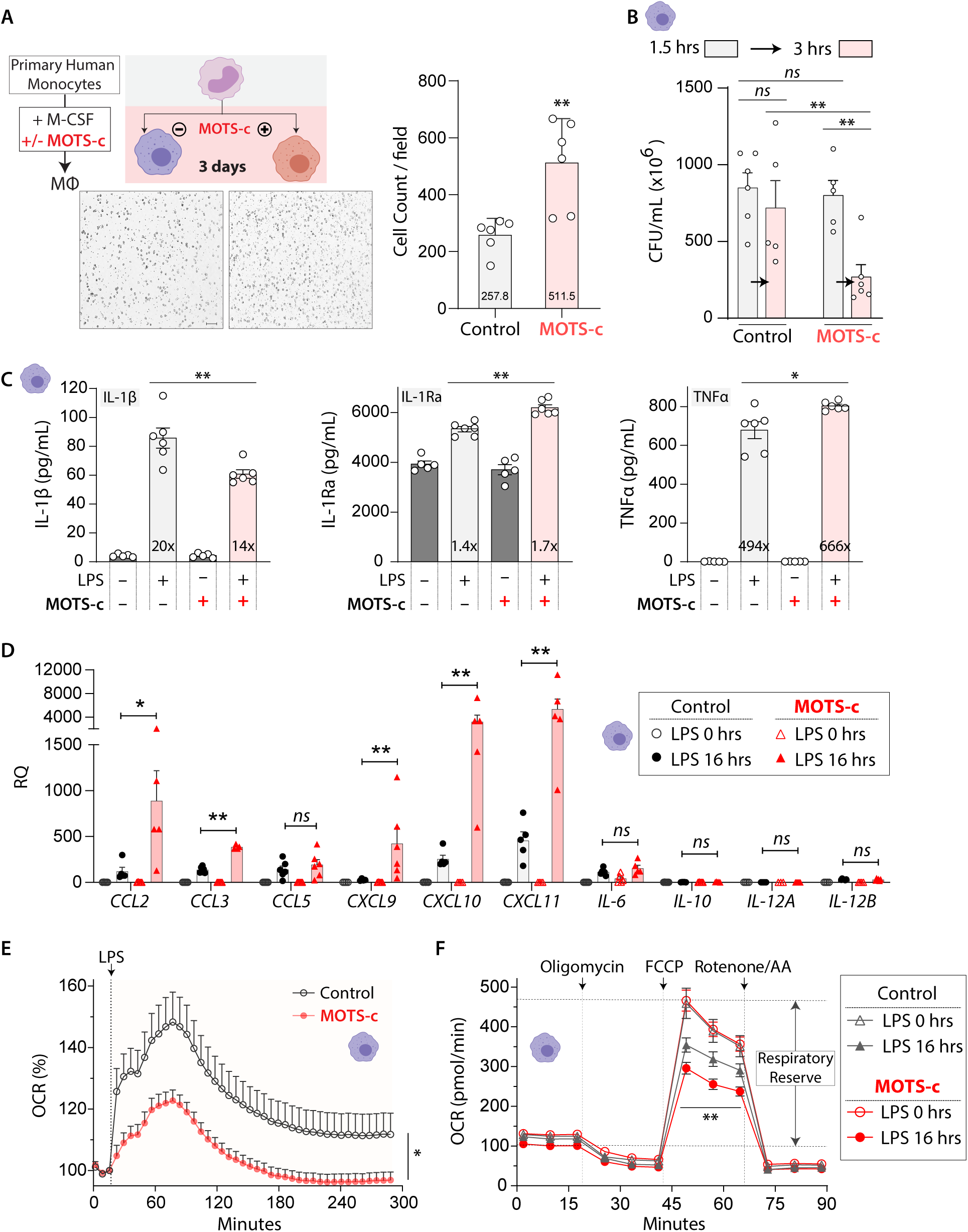
MOTS-c promotes the generation of macrophages with enhanced antibacterial capacity. (**A**) Primary human monocytes were differentiated by M-CSF for 3 days with MOTS-c (10 µM) or vehicle (ddH_2_O)(2-hour priming, then single treatment with M-CSF). Representative images of adhered macrophages (n=6; MΦ=macrophage; Bar, 10 µm). (**B-F**) THP-1 macrophages were differentiated for 4 days with/without MOTS-c treatment (10 µM; 2-hour priming, then single treatment with PMA). (**B**) Gentamicin protection assay in MOTS-c-programmed THP-1 macrophages following 1.5- or 3-hours post-infection of *E. coli* (MOI: 10). CFU: colony forming units (n=6). (**C-F**) MOTS-c-programmed THP-1 macrophages were stimulated with LPS (100 ng/ml) and (**C**) secreted levels of IL-1β, IL-1Ra, and TNFα measured by ELISA after 20 hours (n=6), (**D**) cytokine expression levels determined by RT-qPCR after 16 hours (n=6), and (**E-F**) metabolic flux assessed (n=15) by cellular respiration (oxygen consumption rate; OCR) (**E**) immediately after LPS stimulation and (**F**) 16 hours after LPS stimulation. Data expressed as mean +/- SEM. Mann-Whitney test, except for (E, F), which used two-way ANOVA repeated measures. *P<0.05, ** P<0.01, *** P<0.001.

### Exposure of primary mouse bone marrow cells to MOTS-c promotes the emergence of a distinct subset of mature macrophages in both sexes throughout aging

MOTS-c has a significant impact on aging physiology^19,24^, in part, by acting as a regulator of adaptive stress responses^19,21–24,102,103^ and is considered an emerging mitochondrial hallmark of aging^104,105^. In addition, aging is also accompanied by maladaptive immune responses, including a shift in macrophage function^106–115^. Thus, we tested the impact of early MOTS-c exposure on the differentiation trajectories of primary progenitors from the bone marrow of female and male, young and old C57BL/6JNia mice (**Figure 6A**). Importantly, we used single-cell analyses to investigate heterogeneity in resulting primary macrophage populations (*i.e.* different macrophage “states”), reflecting their broad spectrum of transcriptional programs trained to dynamically adapt to their environment^116–118^.

**Figure 6.**
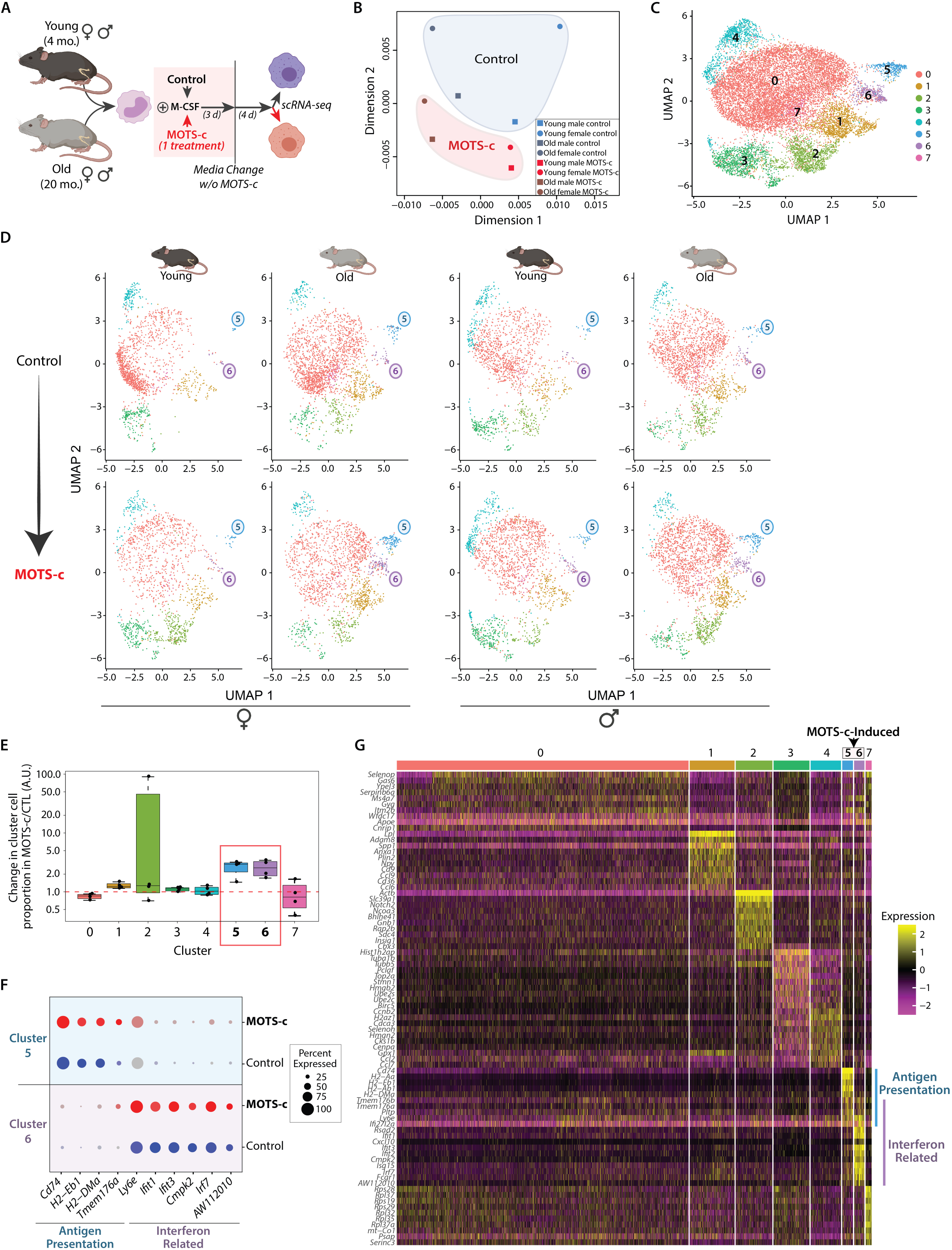
MOTS-c generates unique macrophages characterized by enhanced interferon signaling and antigen presentation in an age-related manner. (**A**) Single-cell RNA-seq (scRNA-seq) was performed on bone marrow-derived macrophages (BMDMs) from young (4 mo.) and old (20 mo.) mice of both sexes that were differentiated for 7 days in the presence of MOTS-c (10 µM) or vehicle (ddH_2_O), treated once concomitantly with first exposure to M-CSF, and present only for the first 3 days of differentiation. (**B**) Multidimensional scaling (MDS) analysis across each of the 8 groups based on pseudobulk gene expression profiles for each biological group after performing DESeq2 VST normalization. (**C-D**) Uniform manifold approximation and projection (UMAP) plot on (**C**) all mice and (**D**) separated by age and sex, with cells color-coded based on SNN-clustering. Clusters 5 and 6 were enriched in MOTS-c-programmed BMDM populations. (**E**) Box plot of relative cluster cell proportion ratios between MOTS-c-programmed *vs.* control macrophages across clusters. Note that clusters 5 and 6 are consistently found in higher proportion in MOTS-c treated samples compared to their corresponding control condition. (**F**) Dotplot of select genes enriched in clusters 5 and 6 (see also **Figures S10-S12** and **Table S3**). (**G**) Heatmap of the top 10 differential gene markers of each of the 8 clusters in BMDMs induced in the presence/absence of MOTS-c (FDR < 5%).

Specifically, we used single-cell RNA-seq (scRNA-seq) to determine whether early exposure (during the first 3 days of differentiation) of primary mouse bone marrow progenitors to MOTS-c may promote the emergence of transcriptionally distinct macrophage subsets. Because of the age-dependent shift in macrophage adaptive capacity^119^, we collected bone marrow cells from young (4 mo.) and old (20 mo.) mice of both sexes and differentiated them for 7 days +/- MOTS-c into bone marrow-derived macrophages (BMDMs). MOTS-c was given only once concomitantly with M-CSF at the onset of differentiation, and the media was replaced after 3 days in both the control- and MOTS-c-treated conditions (**Figure 6A**). scRNA-seq was performed at day 7 of differentiation, generating libraries for each biological group separately (N = 5 animals per group, one library for each group obtained by equicellular mixing of BMDMs from each animal). To minimize the negative impact of batch effects, all samples were processed in parallel from bone marrow collection to library preparation and sequencing.

First, we asked whether there were global changes to mature BMDM transcriptomes upon only early exposure to MOTS-c, irrespective of age and sex. For this purpose, we decided to leverage a pseudobulk approach, which best controls false discovery rates^120^. After aggregating reads for each independent library, we normalized the data using the Variance Stabilizing Transformation from the ‘DESeq2’ R package. Multidimensional scaling (MDS) revealed that macrophages were clearly separated by age on dimension 1 and by MOTS-c treatment on dimension 2 (**Figure 6B**). Intriguingly, female BMDMs showed greater shift in response to MOTS-c treatment than male BMDMs; notably, female MOTS-c-treated BMDMs appeared “masculinized” (*i.e.* closer to male samples in MDS space, and with reduced sample-to-sample distance; **Figures 6B and S9**). With limited sample number, future work will be needed to elucidate interactions of MOTS-c treatment with age and sex in the context of BMDM transcriptional programming. However, our pseudobulk analysis reveals global remodeling of BMDM programs upon transient early exposure to MOTS-c.

Using a shared nearest neighbor (SNN) modularity optimization with the ‘Seurat’ R package, we identified 8 BMDM clusters, indicative of different latent transcriptional states, consistent with a heterogeneous mature macrophage population. Importantly, the 8 distinct single-cell BMDM clusters were comprised of cells from young and old mice of both sexes [cluster labeling is shown on our data using a nonlinear dimensionality-reduction technique, UMAP (uniform manifold approximation and projection)] (**Figure 6C**). Intriguingly, 2 clusters (clusters 5 and 6) were consistently enriched in MOTS-c-treated condition and their increase was more pronounced in BMDMs derived from older animals regardless of sex (**Figure 6D and 6E**; **Table S3**). To note, aging alone increased the proportion of clusters 5 and 6 (**Figure 6D**; upper panels), consistent with the connection between MOTS-c signaling and aging^19,24,121–123^. Cluster 5 was largely enriched in the expression of genes relevant to antigen presentation, whereas cluster 6 showed increased expression of interferon-related genes (**Figures 6F**-**6G and S10-S11**; full analysis in **Table S3**); some genes were shared between these categories. Consistently, overrepresentation analysis using Gene Ontology Biological Process terms (GOBP) showed a clear signature of antigen presentation processes for cluster 5 and IFN-related processes for cluster 6 (FDR < 5%; top 20 pathways in **Figure S12**; full analysis in **Table S4**). Notably, interferon signaling can be triggered not only by mtDNA^2^, but can also induce the expression mtDNA-encoded MOTS-c (**Figure 3C and 3E**)^66^. Increased expression of genes involved in antigen presentation and interferon signaling with age has been observed in monocytes and macrophages in mice^124–128^. In fact, the antigen presentation genes *H2-Aa, H2-Ab1, H2-Eb1, CD74,* and *AW112010* and interferon-related genes *Irf7, Ifit2, Ifit3, Ifitm3*, and *Ifi204* are consistently upregulated with age in our data and that of others (**Table S3**)^124–127^, indicating a conserved effect of aging on monocytes/macrophages.

## Discussion

Here, we identify MOTS-c as a first-in-class mitochondrial-encoded HDP that can target bacteria and modulate monocyte differentiation. MOTS-c influenced nuclear gene expression during the early phase of monocyte-to-macrophage differentiation to generate distinct macrophage populations that are adapted to bacterial clearance. Our current working model is that MOTS-c initially mounts a preemptive attack on bacteria by aggregating them and preventing growth, then influences monocyte differentiation for efficient microbial clearance. Notably, HDPs possess anti-inflammatory effects upon clearing pathogens to promote a non-inflammatory post-infection resolution and alleviate over-responsive inflammation (*e.g.* sepsis)^129,130^, consistent with the systemic anti-inflammatory effects of MOTS-c^19,21,23,31,131–143^.

Our initial hypothesis was that interbacterial communication using peptides may still exist between bacteria-derived mitochondria and bacteria. Endosymbiosis is a form of sustained infection whereby the primal bacteria and our ancestral cell communicated to coordinate the establishment of mitochondria and eukaryotic life. Such communication was likely derived from immunological processes to regulate each other, which may have laid the foundation for cellular signal transduction. HDPs often regulate highly-conserved cellular functions, such as ribosomes and protein metabolism^144^, consistent with the stress-responsive regulatory functions of MOTS-c in mammalian cells^102,103,145–149^(**Figure 4E**-**4H**). Evidence for MOTS-c as a therapeutic agent against MRSA in mice adds translational potential for mtDNA-encoded immune peptides for combating bacterial infection^150^. Further, it is likely that MOTS-c, consistent with other HDPs, may also target viruses. Human myeloblast cells that were infected with Sendai virus to induce IFN expression showed considerable induction of transcripts from the mitochondrial rRNA loci (∼75% of induced transcripts)^66^, a study that influenced the discovery of MOTS-c^19^. Consistently, we found that MOTS-c is endogenously expressed in monocytes and that it can be induced by IFNγ stimulation (**Figure 3E**).

Our data supports an immunological role for mitochondrial-encoded MOTS-c and provides a proof-of-principle for mitochondrial-encoded HDPs. Together with the ever-increasing identification of sORFs in both our mitonuclear genomes, multiple unannotated HDPs may exist in humans with the potential for clinical use against a broad range of infections^151,152^.

## Methods

### Bacterial Strains

*E. coli* [BL21 (DE3); NEB], MRSA (methicillin-resistant *S. aureus*)(ATCC 33592), *S. typhimurium* (ATCC 14028), and *P. aeruginosa* (ATCC 27853) were used. *E.coli* cells were transformed to conditionally overexpress MOTS-c in pSF-T7/LacO (Sigma, OGS500) using IPTG. Bacteria were routinely maintained in LB medium (liquid and agar). A modified M9 minimal medium, composed of 0.1x of M9 Minimal Salts without NaCl (BD Difco), 1 mM MgSO_4_, 1% glucose, and 2.5 g/L peptone was used in select studies.

### Phase-Contrast Light Microscopy

Bacteria were grown overnight in LB broth, from which a 1:100 inoculation was made in LB broth and grown to mid-log phase (∼0.6 OD_600_), measured using a SpectraMax M3 Spectrophotometer (Molecular Devices), in a 37°C shaker at 225 rpm. 1 mL of mid-log *E. coli* (BL21) and 3 mL of mid-log MRSA were collected and treated in modified M9 minimal medium to achieve macroscopic bacterial aggregates. Approximately 3 µL of aggregates were transferred to Superfrost Plus Micro Slides (VWR) with platinum-grade cover slips and imaged with phase-contrast mode using a EVOS FL Cell Imaging System.

### Scanning Electron Microscopy

Samples were fixed overnight at 4 °C in 3% glutaraldehyde. Fixed cells were placed onto 0.2-μm Nuclepore Track-Etch Membrane filters (Whatman) and allowed to air dry for fifteen minutes prior to ethanol dehydration. The dehydration series progressed from an initial wash concentration of 30% ethanol with 30-minute stepwise increments to a final wash of 100% ethanol prior to critical point drying (Autosamdri-815, Toursimis). Samples were then sputter coated (Cressington) with approximately 3 nm of Pd. Electron micrographs were obtained with a JEOL-7001 FEG Scanning Electron Microscope.

### Confocal imaging

Cellular images were obtained using a Zeiss LSM700 confocal microscope system (Germany). THP-1 monocytes were treated with synthetic MOTS-c peptide tagged with a FITC fluorophore (1 μM) for 30 min. After washing 3 times with PBS, cells were fixed using 4% paraformaldehyde and permeabilized in 0.2% TritonX100. Fixed cells were incubated with DAPI (Sigma) for 30 min and washed an additional three times with 0.1% PBST. The cells were spread onto glass coverslips and attached to MicroSlides (cat#48311-703, VWR) using ProLong Gold antifade mountant (cat#P36934, ThermoFisher).

### Bacterial growth measurements

Overnight bacterial cultures were diluted 1:1000 and grown in modified M9 minimal medium (0.1x M9, 5% glucose, 1mM MgSO4, 1.0 g/L peptone) with MOTS-c (or vehicle) in a 37°C shaker at 225 rpm. Bacterial density was measured at OD_600_ every hour in two-sided polystyrene cuvettes (VWR) using a SpectraMax M3 Spectrophotometer (Molecular Device). For plate assays, modified M9 minimal medium agar (1.5%) plates (35 mm Petri dishes) were made.

### ATP Assay

BacTiter-Glo Microbial Cell Viability Kit (Promega) was used to assess bacterial cell viability and metabolism by measuring luminescence correlated with amount of intracellular ATP. Bacteria were grown overnight in LB broth, from which a 1:100 inoculation was grown in LB broth to reach mid-log phase (∼0.6 OD_600_) in a 37°C shaker at 225 rpm. 1 mL of log-phase bacteria was collected, resuspended in modified M9 minimal medium, treated with MOTS-c (100 µM), then subject to luciferase reaction per the manufacturer’s instructions. Luminescence was measured using SpectraMax M3 Spectrophotometer (Molecular Device) and values reported as RLU (relative light units).

### SYTOX Green Assay

SYTOX® Green Nucleic Acid Stain 5 mM in DMSO (ThermoFisher) was used to assess bacterial membrane integrity. SYTOX® Green does not cross intact membranes, but easily penetrates compromised membranes and stains nucleic acids ^52^. Fluorescence readings (bottom-read, excitation/emission at 504/523 nm) were recorded at various time points with one measurement taken before treatment (blank).

### Western Blot

Whole cell and nuclear compartment were lysed with 8 M Urea buffer containing a protease/phosphatase inhibitor cocktail (Thermo Fisher Scientific) and were sonicated for 15 seconds at 60% amplitude. The lysates were separated by 8-16% pre-cast SDS-PAGE gels (Bio-Rad) and transferred onto PVDF membranes. The membranes were blocked with 5% bovine serum albumin (BSA) in tris-buffered saline with 0.05% Tween-20 and probed with primary antibody at 4°C overnight. Proteins of interest were detected with anti-rabbit IgG HRP linked antibody (Cell Signaling) and developed by Clarity Western ECL substrates (Bio-Rad). The membranes were imaged by the ChemiDoc XRS+ system (Bio-Rad).

### Human cell culture

THP-1 cells (RRID:CVCL_0006) were routinely cultured at a range of cell density of 2×10^5^ cells/ml to 8×10^5^ cells/ml in RPMI 1640 (Corning) supplemented with 10% heat-inactivated fetal bovine serum (Omega Scientific) and 0.05 mM 2-Mercaptoethanol at 37°C with 5% CO_2_. Macrophages were generated by culturing THP-1 (6×10^5^/ml) cells with 15 nM phorbol 12-myristate 13-acetate (PMA, Sigma-Aldrich) in the media. Human primary monocytes were isolated from leukocyte cones of healthy blood donors, obtained from USC/CHLA. Mononuclear cells were separated by centrifugation at 400x g for 20 minutes with underlying Histopaque-1077. The opaque interface between plasma and the Histopaque-1077 was collected for further purification by Red Blood Cell lysis buffer (Miltenyi). Monocytes were isolated by negative selection using the StraightFrom LRSC CD14 MicroBead Kit (cat# 130-117-026, Miltenyi). Purified human monocytes were seeded at a density of 1×10^6^ cells/ml in RPMI 1640 supplemented with 10% heat inactivated fetal bovine serum and 100 ng/ml human M-CSF (Miltenyi).

### Human cell treatment

THP-1 cells were seeded at a density of 4×10^5^ cells/ml and reached to 6×10^5^ cells/ml prior to treatment. PMA (Sigma-Aldrich) was reconstituted in DMSO and used at 15 nM. Lipopolysaccharides from Escherichia coli O111:B4 (List Biology Laboratories, Inc.) were reconstituted in sterile water and used at 100 ng/ml. Human interferon-gamma (Sigma) was reconstituted in sterile water and used at 20 ng/ml. Synthetic MOTS-c peptide with a purity >95% by mass spectrometry (New England Peptides, now Biosynth) was reconstituted in sterile water and used at 10 µM.

### Nuclear fractionation

Nuclear fraction was isolated as previously described ^21,153^. Cells were harvested by centrifugation. The pellet was resuspended in hypotonic fractionation buffer (10 mM Tris, pH 7.5, 10 mM NaCl, 3 mM MgCl2 0.3% NP-40, 10% glycerol, and EDTA-free protease inhibitor) and incubated on ice for 30 minutes and passed through a 31-gauge needle 5 times and centrifuged. The nuclear pellet was washed with hypertonic fractionation buffer 3 times and resuspend in 8 M urea lysis buffer (1 M Tris, pH = 8).

### Gentamicin Protection Assay

THP-1 monocytes were seeded on a 6-well plate at concentration of 1.5×10^6^ cells/well and differentiated by PMA (15 nM) for 96 hours. For MOTS-c treatment, monocytes were first primed with MOTS-c (10 µM) for 2 hours, then treated only once with a mixture of PMA (15 nM) and MOTS-c (10 µM). *E. coli* (BL21) were added to differentiated THP-1 cells (MOI 10) and co-cultured for 1-2.5 hours, at which time gentamicin (50 µg/ml) (Fisher Scientific) was added. 30 minutes later, cells were washed with PBS and collected (total of 1.5 or 3 hours). Cells were lysed with 1% Triton X-100 and spread on LB agar plate. Bacterial colonies were counted after overnight incubation at 37°C.

### ELISA

THP-1 monocytes (n=6) were cultured in six-well plates at a density of 5 x 10^6^ cells/mL in 2 mL of RPMI 1640 (cat#45000-396, VWR, USA) supplemented with 10% fetal bovine serum (Omega Scientific) and incubated at 37°C in humidified air with 5% CO_2_. THP-1 cells were primed with MOTS-c peptide (10 μM) for 2 hours, then treated with a combination of PMA (15 nM) + MOTS-c (10 μM). 96 hours post-differentiation, cells were treated with LPS (100 ng/mL) or vehicle for an additional 20 hours and supernatant were collected for cytokine analysis. The production levels of cytokines including IL-1β (cat#BMS224-2), IL-1Ra (cat#BMS2080), and TNF-α (cat#BMS223-4) were measured in duplicate using human ELISA kits following the manufacturer’s instructions (ThermoFisher) and SpectraMax M3 (Molecular devices).

### Real-time qPCR

Total RNA was extracted and purified using the Direct-zol RNA MiniPrep kit (Zymo Research) following manufacturer’s instructions. The total RNA (1.2 μg) was used to synthesize single-stranded cDNA using iScript cDNA synthesis Kit (Bio-Rad) and T100 Thermal Cycler (Bio-Rad) according to the manufacturer’s instructions. The relative mRNA levels of *CCL2, CCL3, CCL5, CXCL9, CXCL10, CXCL11, IL-6, IL-10, IL-12A,* and *IL-12B* were determined by the real-time quantitative PCR analysis using the SYBR Green Supermix (#1725275, Bio-Rad) and CFX Connect Real-time PCR Detection System (Bio-Rad). All reactions were performed in total of 20 μL reaction volume containing 2 μL of diluted cDNA, 10 μL of SYBR Supermix, 1 μL of 10 μM forward primer, 1 μL of 10 μM reverse primer, and 6 μL of autoclaved distilled water. All samples were analyzed in duplicate with an endogenous control gene (β-Actin) being analyzed at the same time. The relative expression of each gene was calculated from 2^-ΔΔCT^.

### Primer sequences used in real-time RT-qPCR

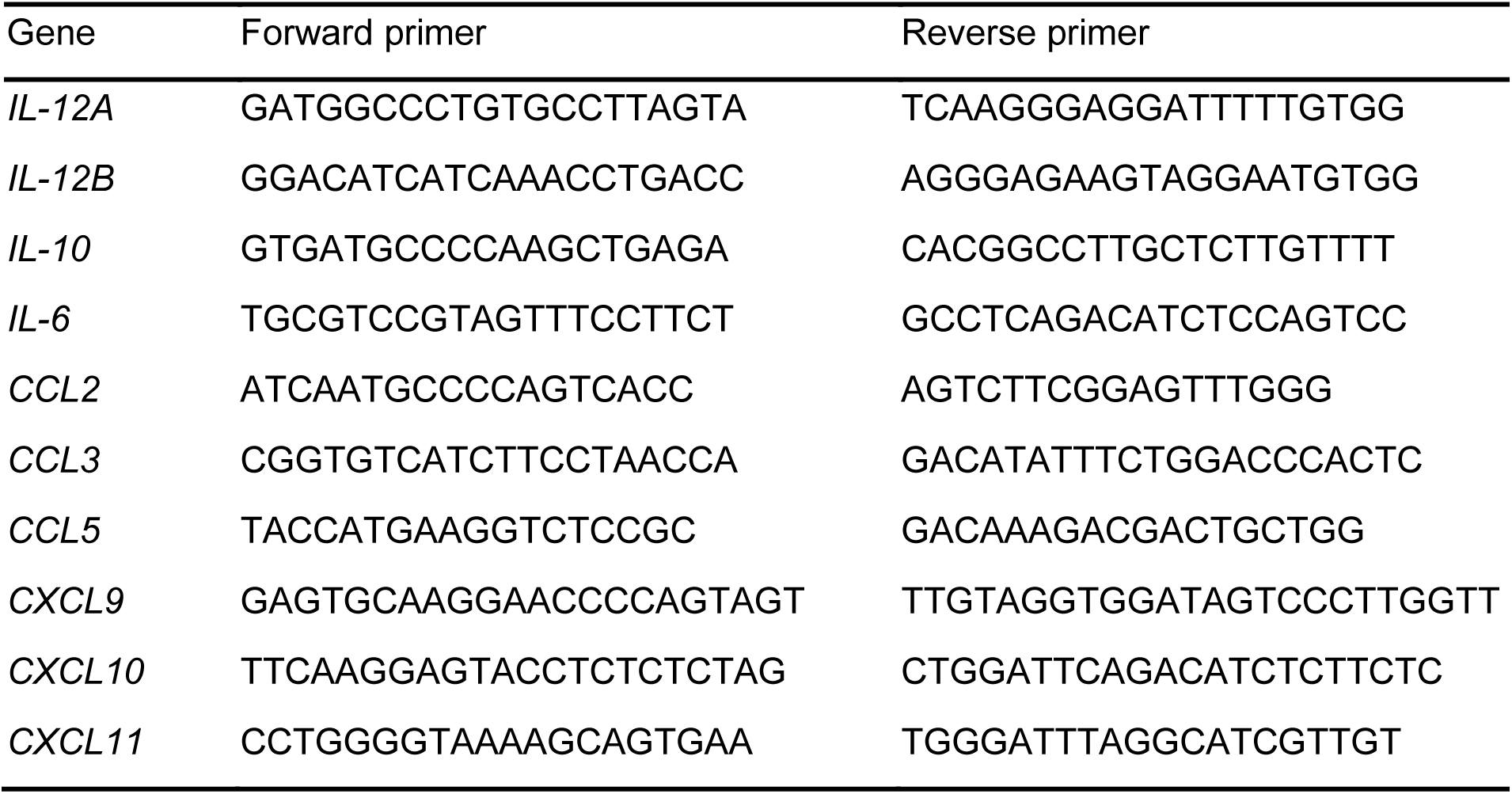

### Metabolic flux measurements

Real-time analysis of oxygen consumption rates (OCR) were measured in THP-1 cells using XF24/96 Extracellular Flux Analyzer (Seahorse Bioscience). Cells were seeded in Seahorse XF96 cell culture microplate (#101085-004, Seahorse Bioscience) at a density of 5 x 10^6^ cells/mL in 100 μL of RPMI 1640 (#45000-396, VWR) supplemented with 10% fetal bovine serum (Omega Scientific). The cells were treated without or with 10 μM MOTS-c for 2 hours, followed by treatment with PMA for 96 hours. LPS (100 ng/mL) was added either during OCR measurements (dispensed by XF96) or added for 16 hours prior to OCR measurements and medium was replaced with XF assay buffer supplemented with 1 mM pyruvate and 12 mM D-glucose. ATP turnover, maximum respiratory capacity, and non-mitochondrial respiration were estimated by sequential addition of oligomycin (0.9 μM), carbonyl cyanide 4-[trifluoromethoxy]phenylhydrazone (FCCP, 1.0 μM), rotenone (0.5 μM), and antimycin A (0.5 μM). All readings were normalized to relative protein concentration.

### Bulk RNA-seq library preparation

1 μg of total RNA was subjected to rRNA depletion using the NEBNext rRNA Depletion Kit (New England Biolabs), according to the manufacturer’s protocol. Strand specific RNA-seq libraries were then constructed using the SMARTer Stranded RNA-Seq Kit (Clontech #634839), according to the manufacturer’s protocol. Based on rRNA-depleted input amount, 13-15 cycles of amplification were performed to generate final RNA-seq libraries. Pooled libraries were sent for paired-end sequencing on the Illumina HiSeq-Xten platform at the Novogene Corporation (USA). The raw sequencing data has been deposited to the NCBI Sequence Read Archive (accession number PRJNA623667). The resulting data was analyzed with a standard RNA-seq data analysis pipeline (described below).

### Bulk RNA-seq analysis pipeline

To avoid the mapping issues due to overlapping sequence segments in paired-end reads, reads were hard trimmed to 75bp using Fastx toolkit v0.0.13. Reads were then further quality-trimmed using Trimgalore 0.4.4 to retain high-quality bases with Phred score > 20. All reads were also trimmed by 6 bp from their 5’ end to avoid poor qualities or random-hexamer driven sequence biases. cDNA sequences of protein coding and lncRNA genes were obtained through ENSEMBL Biomart for the GRCh38 build of the human genome (release v96). Trimmed reads were mapped to this reference using kallisto v0.43.0 and the –fr-stranded option ^154^. All bulk RNA-seq analysis were performed in the R statistical software Version 3.4.1 (https://cran.r-project.org/). Read counts were imported into R, and summarized at the gene level to estimate gene expression levels. We estimated differential gene expression between control, PMA and PMA+MOTS-c treated THP-1 RNA-seq samples using the ‘DESeq2’ R package (DESeq2 1.16.1)^155^. The heatmap of expression across samples for significant genes (**Figure 5D**) was plotted using the R package ‘pheatmap’ 1.0.10 ^156^.

### Functional enrichment analysis

To perform functional enrichment analysis, we used the Gene Set Enrichment Analysis (GSEA) paradigm through R packages ‘phenoTest’ 1.24.0 and ‘qusage’ 2.10.0. Gene Ontology (GO) gene sets were obtained from the Molecular Signature Database, C5 collection (c5.all.v6.2.symbols.gmt)^157^. A False Discovery Rate (FDR) threshold of 0.05 was considered statistically significant. The -log10(FDR) value for GSEA enrichment is reported in Figure 3F as a barplot for selected terms, and FDR values of 0 were replaced by a small value of 10^-30^ to enable plotting on a reasonable scale. The full list of enriched GO terms at FDR 0.05 is reported in **Table S2**.

### STRING analysis

The list of significantly regulated genes in the PMA vs. PMA+MOTS-c conditions at FDR < 0.05 was used to infer potential disruption to protein interaction networks using the STRING (Search Tool for the Retrieval of Interacting Genes/Proteins) tool database version 11.0 ^90^.

### Mouse husbandry

All animals were treated and housed in accordance with the Guide for Care and Use of Laboratory Animals. All experimental procedures were approved by the University of Southern California’s Institutional Animal Care and Use Committee (IACUC) and are in accordance with institutional and national guidelines. For murine peritonitis experiments, male and female C57BL/6J mice were obtained from Jackson Laboratory and experiments performed at 4-6 months of age. For single-cell RNA-seq analyses, male and female C57BL/6JNia mice (4 and 20 month old animals) were obtained from the National Institute on Aging (NIA) colony at Charles Rivers. Animals were acclimated at the SPF animal facility at USC for 2-4 weeks before any processing, and were euthanized between 8-11am to minimize circadian effects. In all cases, animals were euthanized using a “snaking order” across all groups to minimize batch-processing confounds. All animals were euthanized by CO_2_ asphyxiation followed by cervical dislocation.

### Murine model of peritonitis

MRSA (methicillin-resistant *S. aureus*; ATCC 33592) were grown overnight in LB broth, from which a 1:100 inoculation was made in LB broth and grown to mid-log phase (∼0.6 OD_600_), measured using a SpectraMax M3 Spectrophotometer (Molecular Devices), in a 37°C shaker at 225 rpm. 6×10^8^ or 4×10^8^ CFU of MRSA was washed and resuspended in 100 μM MOTS-c or water immediately prior to intraperitoneal injection with a 28G syringe. MRSA was serial diluted, plated on LB agar, and colonies counted after overnight incubation to confirm CFU inoculated. Mice were monitored for 72 hours and weighed daily. Moribund animals were euthanized by CO_2_ asphyxiation followed by cervical dislocation according to IACUC approved experimental guidelines. To heat-kill MRSA, MRSA resuspended in vehicle (water) was heated in a 70°C water bath for 20 minutes. Mice were euthanized either 6 hours or 72 hours post-inoculation and blood collected by cardiac puncture. Plasma was collected and stored at -80°C after centrifugation at 800g for 10 minutes at 4°C. Cytokines and organ injury markers were measured in plasma following manufacturer’s instructions using the Meso Scale Discovery V-PLEX Proinflammatory Panel 1 Mouse Kit (K15048D-1) for cytokines, Mouse ALT ELISA Kit (ab282882), Mouse AST ELISA Kit (ab263882), and Urea Assay Kit (ab83362).

### Derivation of bone marrow-derived macrophages (BMDMs) and MOTS-c treatment

We isolated BMDMs as previously described ^158^. Briefly, the long bones of each mouse were harvested and kept on ice in D-PBS (Corning) supplemented with 1% Penicillin/Streptomycin (Corning) until further processing. Muscle tissue was removed from the bones, and the bone marrow from cleaned bones was collected into clean tubes ^159^. Red blood cells from the marrow were removed using Red Blood Cell Lysis buffer (Miltenyi Biotech #130-094-183), according to the manufacturer’s instructions, albeit with no vortexing step and incubation for only 2 minutes. The suspension was filtered on 70μm mesh filters to retain only single cells. Cells were plated in macrophage growth medium [DMEM/F12 (Corning), 10% FBS (Sigma), 1% Penicillin/Streptomycin (Corning), 2 ng/mL recombinant M-CSF (Miltenyi Biotech), and 10% L929-conditioned medium as an additional source of M-CSF]. For cells undergoing MOTS-c treatment, this initial culture medium was supplemented with 10 μM MOTS-c peptide (New England Peptides, >95% purity). After 3 days, adherent cells were rinsed with D-PBS, and fresh macrophage growth media was added. Cells were collected after 7 days in culture, when BMDMs have completed differentiation. We differentiated BMDMs from 5 animals from each group (young females, young males, old females and old males, control and MOTS-c treated; 8 groups total).

### BMDM single-cell RNA-seq library preparation

Single-cell RNAseq libraries were prepared using Single Cell 3′ v2 Reagent Kits, according to the manufacturer’s instructions (10xGenomics). For single-cell RNA-seq profiling, BMDMs were detached using ice-cold 10 mM EDTA, counted using a COUNTESS cell counter (Thermo Fisher Scientific), and samples from each sample group were pooled in an equicellular mix (8 mixes, 1 mix per biological condition). Using the 10xGenomics Single Cell 3′ v2 manufacturer’s instructions (10xGenomics), we loaded the microfluidics device with a targeted capture in each sample of 3,000 cells. The 8 samples were run in parallel on the same microfluidics chip and processed on a Chromium Controller instrument (10xGenomics), to generate single-cell Gel bead-in-EMulsions (GEMs). GEM-RT was performed in a C1000 Touch Thermal Cycler with a deep well module (Biorad). The cDNA was amplified and cleaned up with SPRIselect Reagent Kit (Beckman Coulter Genomics). Single-indexed sequencing libraries were constructed using Chromium Single-Cell 3′ Library Kit. Library quality and quantification prior to pooling were assessed using a D1000 screentape device on the Tapestation apparatus (Agilent). All barcoded libraries were combined in a single pool for sequencing, and sent for sequencing on 3 lanes of Hiseq-X-Ten at Novogene Corporation as paired end 150bp reads. The final average sequencing depth per cell was ∼70,000 reads per cell.

### BMDM single-cell RNA-seq data analysis

Reads were hard trimmed to yield the lengths expected by the CellRanger pipeline (Read1: 26bp, Read 2: 98bp) using the fastx_trimmer tool from the FASTX Toolkit v0.0.13 [http://hannonlab.cshl.edu/fastx_toolkit/]. The raw sequencing data has been deposited to the NCBI Sequence Read Archive (accession number PRJNA769064). Trimmed reads were then processed using CellRanger software version 3.0.2 and the mm10 mouse genome reference for mapping, cell identification and UMI processing (10X Genomics). Analyses for single-cell RNA-seq were performed using R version 3.6.3 (single-cell level clustering and marker identification) or 4.1.2 (pseudobulk analysis).

Since sequencing on new generation Illumina patterned flow cells can lead to the emergence of non-physiological chimeric reads, phantom molecules were identified and removed from the cellranger h5 output using the PhantomPurge 1.0.0 R package ^160^. After purging, data was converted to the ‘Seurat’ format for single cell analysis for analysis with Seurat 3.2.2 package ^161^. Genes detected in at least 50 cells, and cells with greater than 1000 genes and less than 10% of mitochondrial genes were selected for downstream analysis, yielding 15,415 cells and 11,622 genes passing quality control filters. The impact of number of genes detected per cells, percentage mitochondrial read and cell cycle phase were regressed out as recommended by the Seurat package. We next leverage the DoubletFinder 2.0.3 package ^162^ to identify and filter out likely cell doublets, assuming a 2.3% doublet formation rate based on 10xGenomics estimates when capturing 3,000 cells per samples. After this, 15,060 cells were identified as singlets by DoubletFinder and used for downstream analyses. Data was normalized using the SCtransform framework. We then ran Principal Component Analysis (PCA), dimensionality reduction using UMAP using 30 principal components, and clustering with a resolution set to 0.2, yielding a total of 8 clusters. Cluster markers were identified using the “FindAllMarkers” function with a minimum 25% of cells and a log_2_fold change threshold of 0.25 using the Wilcoxon-test, and a significance threshold of FDR < 0.05.

To analyze the scRNA-seq dataset with a pseudobulk approach, unnormalized counts were aggregated from cells within the same group (8 groups across age, sex, and MOTS-c treatment). Using R version 4.1.2 and the DESeq2 1.34.0 R package, differential transcriptomic profiles between the groups was estimated and visualized with multidimensional scaling (MDS). To estimate the pairwise distance between female and male pseudobulk samples, we used a distance metric based on Spearman Rank Correlation Rho (1-Rho). The distance of each female sample to both male samples was calculated in the control and MOTS-c treated conditions. To determine whether the distance between female and male samples was reduced upon MOTS-c treatment (as hypothesized based on the MDS analysis), we then used a paired one-sided Wilcoxon Rank sum test to compare the distances in control *vs.* MOTS-c treated conditions.

### BMDM single cell Gene Ontology cluster marker data analysis

To analyze which functional categories were enriched in association to BMDMs in clusters 5 and 6, whose frequency increased upon MOTS-c treatment, we used overrepresentation analysis with clusterProfiler 3.14.3 and annotation package org.Mm.eg.db 3.10.0. Enrichment was computed against the background of all expressed genes detected in the single cell dataset.

## Code availability

The analytical code for the RNA-seq dataset is available on the Benayoun lab github (https://github.com/BenayounLaboratory/MOTSc_Macrophage_Immunity). R code was run using R version 3.4.1, 3.6.3 or 4.1.2 as indicated in relevant sections.

## Acknowledgments

We thank the USC Leonard Davis School of Gerontology Seahorse Bioanalyzer and Aging Biomarker and Service Cores, the USC Translational Imaging Center, the USC Core Center of Excellence in Nano Imaging, and the USC Genomics Core for experimental assistance.

Funding was provided by the NIA (T32 AG052374, F31 AG082606) and an AFAR Diana Jacobs Kalman/AFAR Scholarships for Research in the Biology of Aging to M.C.R., the NIA (T32 AG052374) and an AFAR Diana Jacobs Kalman/AFAR Scholarships for Research in the Biology of Aging to R.J.L., AADOCR Student Research Fellowship awarded to J.S.K., the Larry L. Hillblom Foundation (LLHF) fellowship grant to J.M.S., the NIA (R01 AG076433), Pew Biomedical Scholar award #00034120, and the Kathleen Gilmore Biology of Aging research award to B.A.B., and the NIA (R01 AG052558, R56 AG069955), NIGMS (R01 GM136837), Hevolution, Ellison Medical Foundation (EMF), AFAR, and the Hanson-Thorell Family to C.L.

## Author contributions

B.A.B. and C.L. conceived the experiments. M.C.R., J.S.K., M.I., S.W.J., C.Y.P., R.W.L., C.R.B., J.M.S., K.T., E.K., R.J.L., I.C., B.A.B., and C.L. performed experiments and analyzed the data. C.L. wrote the manuscript with input from authors. All authors approved the manuscript.

## References

1 Martijn, J., Vosseberg, J., Guy, L., Offre, P. & Ettema, T. J. G. Deep mitochondrial origin outside the sampled alphaproteobacteria. Nature 557, 101–105 (2018). 10.1038/s41586-018-0059-5

2 West, A. P. & Shadel, G. S. Mitochondrial DNA in innate immune responses and inflammatory pathology. Nature Reviews Immunology 17, 363–375 (2017). 10.1038/nri.2017.21

3 Papayannopoulos, V. Neutrophil extracellular traps in immunity and disease. Nature Reviews Immunology 18, 134–147 (2018). 10.1038/nri.2017.105

4 Yousefi, S. et al. Catapult-like release of mitochondrial DNA by eosinophils contributes to antibacterial defense. Nature medicine 14, 949–953 (2008). 10.1038/nm.1855

5 Ingelsson, B. et al. Lymphocytes eject interferogenic mitochondrial DNA webs in response to CpG and non-CpG oligodeoxynucleotides of class C. Proceedings of the National Academy of Sciences 115, E478–E487 (2018). 10.1073/pnas.1711950115

6 Chen, J. et al. Pervasive functional translation of noncanonical human open reading frames. Science 367, 1140–1146 (2020). 10.1126/science.aay0262

7 Saghatelian, A. & Couso, J. P. Discovery and characterization of smORF-encoded bioactive polypeptides. Nature chemical biology 11, 909–916 (2015). 10.1038/nchembio.1964

8 Lee, C., Yen, K. & Cohen, P. Humanin: a harbinger of mitochondrial-derived peptides? Trends in endocrinology and metabolism: TEM (2013). 10.1016/j.tem.2013.01.005

9 Kim, S.-J., Xiao, J., Wan, J., Cohen, P. & Yen, K. Mitochondrially derived peptides as novel regulators of metabolism. The Journal of physiology 595, 6613–6621 (2017). 10.1113/jp274472

10 Hancock, R. E., Haney, E. F. & Gill, E. E. The immunology of host defence peptides: beyond antimicrobial activity. Nature reviews. Immunology 16, 321–334 (2016). 10.1038/nri.2016.29

11 Mookherjee, N., Anderson, M. A., Haagsman, H. P. & Davidson, D. J. Antimicrobial host defence peptides: functions and clinical potential. Nature Reviews Drug Discovery (2020). 10.1038/s41573-019-0058-8

12 Zhang, L. J. & Gallo, R. L. Antimicrobial peptides. Current biology : CB 26, R14–19 (2016). 10.1016/j.cub.2015.11.017

13 Lewis, K. Platforms for antibiotic discovery. Nature reviews. Drug discovery 12, 371–387 (2013). 10.1038/nrd3975

14 Zasloff, M. Antimicrobial peptides of multicellular organisms. Nature 415, 389–395 (2002). 10.1038/415389a

15 Hanson, M. A. et al. Synergy and remarkable specificity of antimicrobial peptides in vivo using a systematic knockout approach. eLife 8 (2019). 10.7554/elife.44341

16 Sassone-Corsi, M. et al. Microcins mediate competition among Enterobacteriaceae in the inflamed gut. Nature 540, 280–283 (2016). 10.1038/nature20557

17 Thomas, S., Karnik, S., Barai, R. S., Jayaraman, V. K. & Idicula-Thomas, S. CAMP: a useful resource for research on antimicrobial peptides. Nucleic acids research 38, D774–D780 (2010). 10.1093/nar/gkp1021

18 Lazzaro, B. P., Zasloff, M. & Rolff, J. Antimicrobial peptides: Application informed by evolution. Science 368, eaau5480 (2020). doi:10.1126/science.aau5480

19 Lee, C. et al. The mitochondrial-derived peptide MOTS-c promotes metabolic homeostasis and reduces obesity and insulin resistance. Cell metabolism 21, 443–454 (2015). 10.1016/j.cmet.2015.02.009

20 Lopez-Otin, C., Blasco, M. A., Partridge, L., Serrano, M. & Kroemer, G. Hallmarks of aging: An expanding universe. Cell (2022). 10.1016/j.cell.2022.11.001

21 Kim, K. H., Son, J. M., Benayoun, B. A. & Lee, C. The Mitochondrial-Encoded Peptide MOTS-c Translocates to the Nucleus to Regulate Nuclear Gene Expression in Response to Metabolic Stress. Cell metabolism 28, 516–524 e517 (2018). 10.1016/j.cmet.2018.06.008

22 Kang, G. M. et al. Mitohormesis in Hypothalamic POMC Neurons Mediates Regular Exercise-Induced High-Turnover Metabolism. Cell metabolism 33, 334–349.e336 (2021). 10.1016/j.cmet.2021.01.003

23 Kong, B. S., Min, S. H., Lee, C. & Cho, Y. M. Mitochondrial-encoded MOTS-c prevents pancreatic islet destruction in autoimmune diabetes. Cell reports 36, 109447 (2021). 10.1016/j.celrep.2021.109447

24 Reynolds, J. C. et al. MOTS-c is an exercise-induced mitochondrial-encoded regulator of age-dependent physical decline and muscle homeostasis. Nature communications 12 (2021). 10.1038/s41467-020-20790-0

25 Le, C.-F., Fang, C.-M. & Sekaran, S. D. Intracellular Targeting Mechanisms by Antimicrobial Peptides. Antimicrobial Agents and Chemotherapy 61, AAC.02340-02316 (2017). 10.1128/aac.02340-16

26 Monnet, V., Juillard, V. & Gardan, R. Peptide conversations in Gram-positive bacteria. 1–13 (2014). 10.3109/1040841x.2014.948804

27 Keller, L. & Surette, M. G. Communication in bacteria: an ecological and evolutionary perspective. Nature reviews. Microbiology 4, 249–258 (2006). 10.1038/nrmicro1383

28 Bassler, B. L. Small talk. Cell-to-cell communication in bacteria. Cell 109, 421–424 (2002).

29 Cotter, P. D., Ross, R. P. & Hill, C. Bacteriocins — a viable alternative to antibiotics? Nature Reviews Microbiology 11, 95–105 (2013). 10.1038/nrmicro2937

30 Miller, B., Kim, S.-J., Kumagai, H., Yen, K. & Cohen, P. Mitochondria-derived peptides in aging and healthspan. Journal of Clinical Investigation 132 (2022). 10.1172/jci158449

31 Reynolds, J. et al. MOTS-c is an Exercise-Induced Mitochondrial-Encoded Regulator of Age-Dependent Physical Decline and Muscle Homeostasis. bioRxiv, 2019.2012.2022.886432 (2019). 10.1101/2019.12.22.886432

32 Kyte, J. & Doolittle, R. F. A simple method for displaying the hydropathic character of a protein. Journal of molecular biology 157, 105–132 (1982). 10.1016/0022-2836(82)90515-0

33 Sweet, R. M. & Eisenberg, D. Correlation of sequence hydrophobicities measures similarity in three-dimensional protein structure. Journal of molecular biology 171, 479–488 (1983). 10.1016/0022-2836(83)90041-4

34 Andersson, E. et al. Antimicrobial activities of heparin-binding peptides. 271, 1219–1226 (2004). 10.1111/j.1432-1033.2004.04035.x

35 Chang, R., Subramanian, K., Wang, M. & Webster, T. J. Enhanced Antibacterial Properties of Self-Assembling Peptide Amphiphiles Functionalized with Heparin-Binding Cardin-Motifs. ACS Applied Materials & Interfaces 9, 22350–22360 (2017). 10.1021/acsami.7b07506

36 Melo, M. N., Ferre, R. & Castanho, M. A. Antimicrobial peptides: linking partition, activity and high membrane-bound concentrations. Nature reviews. Microbiology 7, 245–250 (2009). 10.1038/nrmicro2095

37 Sinha, S., Zheng, L., Mu, Y., Ng, W. J. & Bhattacharjya, S. Structure and Interactions of A Host Defense Antimicrobial Peptide Thanatin in Lipopolysaccharide Micelles Reveal Mechanism of Bacterial Cell Agglutination. Scientific reports 7 (2017). 10.1038/s41598-017-18102-6

38 Pulido, D. et al. Antimicrobial Action and Cell Agglutination by the Eosinophil Cationic Protein Are Modulated by the Cell Wall Lipopolysaccharide Structure. Antimicrobial Agents and Chemotherapy 56, 2378–2385 (2012). 10.1128/aac.06107-11

39 Robert, É. et al. Mimicking and Understanding the Agglutination Effect of the Antimicrobial Peptide Thanatin Using Model Phospholipid Vesicles. 54, 3932–3941 (2015). 10.1021/acs.biochem.5b00442

40 Torrent, M., Pulido, D., Nogués, M. V. & Boix, E. Exploring New Biological Functions of Amyloids: Bacteria Cell Agglutination Mediated by Host Protein Aggregation. 8, e1003005 (2012). 10.1371/journal.ppat.1003005

41 Chairatana, P. & Nolan, E. M. Molecular Basis for Self-Assembly of a Human Host-Defense Peptide That Entraps Bacterial Pathogens. 136, 13267–13276 (2014). 10.1021/ja5057906

42 Bahar, A. & Ren, D. Antimicrobial Peptides. Pharmaceuticals 6, 1543–1575 (2013). 10.3390/ph6121543

43 Bals, R., Wang, X., Zasloff, M. & Wilson, J. M. The peptide antibiotic LL-37/hCAP-18 is expressed in epithelia of the human lung where it has broad antimicrobial activity at the airway surface. Proceedings of the National Academy of Sciences of the United States of America 95, 9541–9546 (1998).

44 Kandasamy, S. K. & Larson, R. G. Effect of salt on the interactions of antimicrobial peptides with zwitterionic lipid bilayers. 1758, 1274–1284 (2006). 10.1016/j.bbamem.2006.02.030

45 Choi, H., Rangarajan, N. & Weisshaar, J. C. Lights, Camera, Action! Antimicrobial Peptide Mechanisms Imaged in Space and Time. Trends in Microbiology 24, 111–122 (2016). 10.1016/j.tim.2015.11.004

46 Schmidt, N. W. et al. Criterion for amino acid composition of defensins and antimicrobial peptides based on geometry of membrane destabilization. Journal of the American Chemical Society 133, 6720–6727 (2011). 10.1021/ja200079a

47 Schmidt, N. W. & Wong, G. C. Antimicrobial peptides and induced membrane curvature: geometry, coordination chemistry, and molecular engineering. Curr Opin Solid State Mater Sci 17, 151–163 (2013). 10.1016/j.cossms.2013.09.004

48 Farkas, A., Maroti, G., Kereszt, A. & Kondorosi, E. Comparative Analysis of the Bacterial Membrane Disruption Effect of Two Natural Plant Antimicrobial Peptides. Frontiers in microbiology 8, 51 (2017). 10.3389/fmicb.2017.00051

49 Torrent, M., Pulido, D., Nogues, M. V. & Boix, E. Exploring new biological functions of amyloids: bacteria cell agglutination mediated by host protein aggregation. PLoS Pathog 8, e1003005 (2012). 10.1371/journal.ppat.1003005

50 Sinha, S., Zheng, L., Mu, Y., Ng, W. J. & Bhattacharjya, S. Structure and Interactions of A Host Defense Antimicrobial Peptide Thanatin in Lipopolysaccharide Micelles Reveal Mechanism of Bacterial Cell Agglutination. Scientific reports 7, 17795 (2017). 10.1038/s41598-017-18102-6

51 Robert, E. et al. Mimicking and Understanding the Agglutination Effect of the Antimicrobial Peptide Thanatin Using Model Phospholipid Vesicles. Biochemistry 54, 3932–3941 (2015). 10.1021/acs.biochem.5b00442

52 Roth, B. L., Poot, M., Yue, S. T. & Millard, P. J. Bacterial viability and antibiotic susceptibility testing with SYTOX green nucleic acid stain. Applied and environmental microbiology 63, 2421–2431 (1997). 10.1128/aem.63.6.2421-2431.1997

53 Hurdle, J. G., O’Neill, A. J., Chopra, I. & Lee, R. E. Targeting bacterial membrane function: an underexploited mechanism for treating persistent infections. Nature Reviews Microbiology 9, 62–75 (2011). 10.1038/nrmicro2474

54 Lai, Y. & Gallo, R. L. AMPed up immunity: how antimicrobial peptides have multiple roles in immune defense. Trends in immunology 30, 131–141 (2009). 10.1016/j.it.2008.12.003

55 Sorensen, O., Cowland, J. B., Askaa, J. & Borregaard, N. An ELISA for hCAP-18, the cathelicidin present in human neutrophils and plasma. J Immunol Methods 206, 53–59 (1997).

56 Sorensen, O. E. Human cathelicidin, hCAP-18, is processed to the antimicrobial peptide LL-37 by extracellular cleavage with proteinase 3. Blood 97, 3951–3959 (2001). 10.1182/blood.v97.12.3951

57 Duplantier, A. J. & Van Hoek, M. L. The Human Cathelicidin Antimicrobial Peptide LL-37 as a Potential Treatment for Polymicrobial Infected Wounds. Frontiers in immunology 4 (2013). 10.3389/fimmu.2013.00143

58 Ashrafi, N., Lapthorn, C., Pullen, F. S., Naclerio, F. & Birthe. High performance liquid chromatography coupled to electrospray ionisation mass spectrometry method for the detection of salivary human neutrophil alpha defensins HNP1, HNP2, HNP3 and HNP4. Analytical Methods 9, 6482–6490 (2017). 10.1039/c7ay01676j

59 Lehrer, R. I. & Ganz, T. Defensins: endogenous antibiotic peptides from human leukocytes. Ciba Found Symp 171, 276–290; discussion 290-273 (1992).

60 Nicolas, P. Multifunctional host defense peptides: intracellular-targeting antimicrobial peptides. FEBS Journal 276, 6483–6496 (2009). 10.1111/j.1742-4658.2009.07359.x

61 Cudic, M. & Otvos, L., Jr. Intracellular targets of antibacterial peptides. Current drug targets 3, 101–106 (2002). 10.2174/1389450024605445

62 Shah, P., Hsiao, F. S.-H., Ho, Y.-H. & Chen, C.-S. The proteome targets of intracellular targeting antimicrobial peptides. n/a-n/a (2015). 10.1002/pmic.201500380

63 Ho, Y.-H., Shah, P., Chen, Y.-W. & Chen, C.-S. Systematic analysis of intracellular-targeting antimicrobial peptides, bactenecin 7, hybrid of pleurocidin and dermaseptin, proline-arginine-rich peptide, and lactoferricin B, by using Escherichia coli proteome microarrays. mcp.M115.054999 (2016). 10.1074/mcp.m115.054999

64 Liao, M. H., Chen, S. J., Tsao, C. M., Shih, C. C. & Wu, C. C. Possible biomarkers of early mortality in peritonitis-induced sepsis rats. The Journal of surgical research 183, 362–370 (2013). 10.1016/j.jss.2013.01.022

65 Malanovic, N. & Lohner, K. Gram-positive bacterial cell envelopes: The impact on the activity of antimicrobial peptides. Biochimica et biophysica acta 1858, 936–946 (2016). 10.1016/j.bbamem.2015.11.004

66 Tsuzuki, T. et al. The majority of cDNA clones with strong positive signals for the interferon-induction-specific sequences resemble mitochondrial ribosomal RNA genes. Biochemical and biophysical research communications 114, 670–676 (1983).

67 Auwerx, J. The human leukemia cell line, THP-1: A multifacetted model for the study of monocyte-macrophage differentiation. Experientia 47, 22–31 (1991). 10.1007/bf02041244

68 Chow, N. A., Jasenosky, L. D. & Goldfeld, A. E. A distal locus element mediates IFN-gamma priming of lipopolysaccharide-stimulated TNF gene expression. Cell reports 9, 1718–1728 (2014). 10.1016/j.celrep.2014.11.011

69 Tamai, R., Sugawara, S., Takeuchi, O., Akira, S. & Takada, H. Synergistic effects of lipopolysaccharide and interferon-gamma in inducing interleukin-8 production in human monocytic THP-1 cells is accompanied by up-regulation of CD14, Toll-like receptor 4, MD-2 and MyD88 expression. J Endotoxin Res 9, 145–153 (2003). 10.1179/096805103125001540

70 Duits, L. A., Ravensbergen, B., Rademaker, M., Hiemstra, P. S. & Nibbering, P. H. Expression of beta-defensin 1 and 2 mRNA by human monocytes, macrophages and dendritic cells. 106, 517–525 (2002). 10.1046/j.1365-2567.2002.01430.x

71 Muller, E. et al. Toll-Like Receptor Ligands and Interferon-gamma Synergize for Induction of Antitumor M1 Macrophages. Frontiers in immunology 8, 1383 (2017). 10.3389/fimmu.2017.01383

72 Held, T. K., Weihua, X., Yuan, L., Kalvakolanu, D. V. & Cross, A. S. Gamma interferon augments macrophage activation by lipopolysaccharide by two distinct mechanisms, at the signal transduction level and via an autocrine mechanism involving tumor necrosis factor alpha and interleukin-1. Infect Immun 67, 206–212 (1999).

73 Schroder, K., Hertzog, P. J., Ravasi, T. & Hume, D. A. Interferon-γ: an overview of signals, mechanisms and functions. Journal of leukocyte biology 75, 163–189 (2004). 10.1189/jlb.0603252

74 Nakagomi, A., Freedman, S. B. & Geczy, C. L. Interferon-γ and Lipopolysaccharide Potentiate Monocyte Tissue Factor Induction by C-Reactive Protein. Circulation 101, 1785–1791 (2000). 10.1161/01.cir.101.15.1785

75 Fang, X. M. et al. Differential expression of alpha- and beta-defensins in human peripheral blood. 33, 82–87 (2003). 10.1046/j.1365-2362.2003.01076.x

76 Pioli, P. A. et al. Lipopolysaccharide-Induced IL-1 Production by Human Uterine Macrophages Up-Regulates Uterine Epithelial Cell Expression of Human -Defensin 2. 176, 6647–6655 (2006). 10.4049/jimmunol.176.11.6647

77 Liu, L., Roberts, A. A. & Ganz, T. By IL-1 Signaling, Monocyte-Derived Cells Dramatically Enhance the Epidermal Antimicrobial Response to Lipopolysaccharide. 170, 575–580 (2003). 10.4049/jimmunol.170.1.575

78 Tsutsumi-Ishii, Y. & Nagaoka, I. Modulation of Human -Defensin-2 Transcription in Pulmonary Epithelial Cells by Lipopolysaccharide-Stimulated Mononuclear Phagocytes Via Proinflammatory Cytokine Production. 170, 4226–4236 (2003). 10.4049/jimmunol.170.8.4226

79 Ayabe, T. et al. Secretion of microbicidal α-defensins by intestinal Paneth cells in response to bacteria. Nature Immunology 1, 113–118 (2000). 10.1038/77783

80 Gläser, R. et al. Antimicrobial psoriasin (S100A7) protects human skin from Escherichia coli infection. 6, 57–64 (2005). 10.1038/ni1142

81 Harder, J., Bartels, J., Christophers, E. & Schröder, J. M. A peptide antibiotic from human skin. Nature 387, 861–861 (1997). 10.1038/43088

82 Sun, J. et al. Pancreatic β-Cells Limit Autoimmune Diabetes via an Immunoregulatory Antimicrobial Peptide Expressed under the Influence of the Gut Microbiota. 43, 304–317 (2015). 10.1016/j.immuni.2015.07.013

83 Zhai, Z., Boquete, J.-P. & Lemaitre, B. Cell-Specific Imd-NF-κB Responses Enable Simultaneous Antibacterial Immunity and Intestinal Epithelial Cell Shedding upon Bacterial Infection. Immunity 48, 897–910.e897 (2018). 10.1016/j.immuni.2018.04.010

84 Pena, O. M. et al. Synthetic Cationic Peptide IDR-1018 Modulates Human Macrophage Differentiation. PloS one 8, e52449 (2013). 10.1371/journal.pone.0052449

85 Bowdish, D. M. E., Davidson, D. J., Speert, D. P. & Hancock, R. E. W. The Human Cationic Peptide LL-37 Induces Activation of the Extracellular Signal-Regulated Kinase and p38 Kinase Pathways in Primary Human Monocytes. The Journal of Immunology 172, 3758–3765 (2004). 10.4049/jimmunol.172.6.3758

86 Davidson, D. J. et al. The Cationic Antimicrobial Peptide LL-37 Modulates Dendritic Cell Differentiation and Dendritic Cell-Induced T Cell Polarization. The Journal of Immunology 172, 1146–1156 (2004). 10.4049/jimmunol.172.2.1146

87 Cecotto, L. et al. Antibacterial and anti-inflammatory properties of host defense peptides against Staphylococcus aureus. iScience 25, 105211 (2022). 10.1016/j.isci.2022.105211

88 Drin, G., Cottin, S., Blanc, E., Rees, A. R. & Temsamani, J. Studies on the Internalization Mechanism of Cationic Cell-penetrating Peptides. 278, 31192–31201 (2003). 10.1074/jbc.m303938200

89 Chen, Y. & Meltzer, P. S. Gene Expression Analysis via Multidimensional Scaling. Current Protocols in Bioinformatics 10 (2005). 10.1002/0471250953.bi0711s10

90 Szklarczyk, D. et al. STRING v11: protein–protein association networks with increased coverage, supporting functional discovery in genome-wide experimental datasets. Nucleic acids research 47, D607–D613 (2019). 10.1093/nar/gky1131

91 Khajuria, R. K. et al. Ribosome Levels Selectively Regulate Translation and Lineage Commitment in Human Hematopoiesis. Cell 173, 90–103.e119 (2018). 10.1016/j.cell.2018.02.036

92 Buszczak, M., Robert & Sean. Cellular Differences in Protein Synthesis Regulate Tissue Homeostasis. Cell 159, 242–251 (2014). 10.1016/j.cell.2014.09.016

93 Nicholas, Liana & Jonathan. Ribosome Profiling of Mouse Embryonic Stem Cells Reveals the Complexity and Dynamics of Mammalian Proteomes. Cell 147, 789–802 (2011). 10.1016/j.cell.2011.10.002

94 Shi, Z. & Barna, M. Translating the genome in time and space: specialized ribosomes, RNA regulons, and RNA-binding proteins. Annual review of cell and developmental biology 31, 31–54 (2015). 10.1146/annurev-cellbio-100814-125346

95 Otto, N. A. et al. Adherence Affects Monocyte Innate Immune Function and Metabolic Reprogramming after Lipopolysaccharide Stimulation In Vitro. The Journal of Immunology 206, 827–838 (2021). 10.4049/jimmunol.2000702

96 Viola, A., Munari, F., Sánchez-Rodríguez, R., Scolaro, T. & Castegna, A. The Metabolic Signature of Macrophage Responses. Frontiers in immunology 10 (2019). 10.3389/fimmu.2019.01462

97 Russell, D. G., Huang, L. & Vanderven, B. C. Immunometabolism at the interface between macrophages and pathogens. Nature Reviews Immunology 19, 291–304 (2019). 10.1038/s41577-019-0124-9

98 Langston, P. K., Shibata, M. & Horng, T. Metabolism Supports Macrophage Activation. 8 (2017). 10.3389/fimmu.2017.00061

99 Odegaard, J. I. & Chawla, A. Alternative Macrophage Activation and Metabolism. Annual Review of Pathology: Mechanisms of Disease 6, 275–297 (2011). 10.1146/annurev-pathol-011110-130138

100 Biswas, S. K. & Mantovani, A. Orchestration of Metabolism by Macrophages. Cell metabolism 15, 432–437 (2012). 10.1016/j.cmet.2011.11.013

101 Van Der Does, A. M. et al. LL-37 Directs Macrophage Differentiation toward Macrophages with a Proinflammatory Signature. 185, 1442–1449 (2010). 10.4049/jimmunol.1000376

102 Mottis, A., Herzig, S. & Auwerx, J. Mitocellular communication: Shaping health and disease. Science 366, 827–832 (2019). 10.1126/science.aax3768

103 Galluzzi, L., Yamazaki, T. & Kroemer, G. Linking cellular stress responses to systemic homeostasis. Nature Reviews Molecular Cell Biology 19, 731–745 (2018). 10.1038/s41580-018-0068-0

104 Lopez-Otin, C., Blasco, M. A., Partridge, L., Serrano, M. & Kroemer, G. Hallmarks of aging: An expanding universe. Cell 186, 243–278 (2023). 10.1016/j.cell.2022.11.001

105 Lopez-Otin, C., Galluzzi, L., Freije, J. M., Madeo, F. & Kroemer, G. Metabolic Control of Longevity. Cell 166, 802–821 (2016). 10.1016/j.cell.2016.07.031

106 Franceschi, C., Garagnani, P., Parini, P., Giuliani, C. & Santoro, A. Inflammaging: a new immune–metabolic viewpoint for age-related diseases. Nature Reviews Endocrinology 14, 576–590 (2018). 10.1038/s41574-018-0059-4

107 Pawelec, G. Does the human immune system ever really become ?senescent?? [version 1; referees: 5 approved]. F1000Research 6 (2017). 10.12688/f1000research.11297.1

108 Pawelec, G. Age and immunity: What is “immunosenescence”? Experimental gerontology 105, 4–9 (2018). 10.1016/j.exger.2017.10.024

109 Panda, A. et al. Human innate immunosenescence: causes and consequences for immunity in old age. Trends in immunology 30, 325–333 (2009). 10.1016/j.it.2009.05.004

110 Goronzy, J. J. & Weyand, C. M. Understanding immunosenescence to improve responses to vaccines. Nature Immunology 14, 428–436 (2013). 10.1038/ni.2588

111 Mahbub, S., Deburghgraeve, C. R. & Kovacs, E. J. Advanced age impairs macrophage polarization. Journal of interferon & cytokine research : the official journal of the International Society for Interferon and Cytokine Research 32, 18–26 (2012). 10.1089/jir.2011.0058

112 Lloberas, J. & Celada, A. Effect of aging on macrophage function. Experimental gerontology 37, 1325–1331 (2002). 10.1016/s0531-5565(02)00125-0

113 Plowden, J., Renshaw-Hoelscher, M., Engleman, C., Katz, J. & Sambhara, S. Innate immunity in aging: impact on macrophage function. Aging cell 3, 161–167 (2004). 10.1111/j.1474-9728.2004.00102.x

114 Fei, F., Lee, K. M., McCarry, B. E. & Bowdish, D. M. Age-associated metabolic dysregulation in bone marrow-derived macrophages stimulated with lipopolysaccharide. Scientific reports 6, 22637 (2016). 10.1038/srep22637

115 Minhas, P. S. et al. Macrophage de novo NAD+ synthesis specifies immune function in aging and inflammation. Nature Immunology 20, 50–63 (2019). 10.1038/s41590-018-0255-3

116 Murray, P. J. Macrophage Polarization. Annual review of physiology 79, 541–566 (2017). 10.1146/annurev-physiol-022516-034339

117 Ginhoux, F., Schultze, J. L., Murray, P. J., Ochando, J. & Biswas, S. K. New insights into the multidimensional concept of macrophage ontogeny, activation and function. Nature Immunology 17, 34–40 (2016). 10.1038/ni.3324

118 Xue, J. et al. Transcriptome-Based Network Analysis Reveals a Spectrum Model of Human Macrophage Activation. Immunity 40, 274–288 (2014). 10.1016/j.immuni.2014.01.006

119 Van Beek, A. A., Van Den Bossche, J., Mastroberardino, P. G., De Winther, M. P. J. & Leenen, P. J. M. Metabolic Alterations in Aging Macrophages: Ingredients for Inflammaging? Trends in immunology 40, 113–127 (2019). 10.1016/j.it.2018.12.007

120 Squair, J. W. et al. Confronting false discoveries in single-cell differential expression. Nature communications 12, 5692 (2021). 10.1038/s41467-021-25960-2

121 Fuku, N. et al. The mitochondrial-derived peptide MOTS-c: a player in exceptional longevity? Aging cell 14, 921–923 (2015). 10.1111/acel.12389

122 D’Souza, R. F. et al. Increased expression of the mitochondrial derived peptide, MOTS-c, in skeletal muscle of healthy aging men is associated with myofiber composition. Aging 12 (2020). 10.18632/aging.102944

123 Zempo, H. et al. A pro-diabetogenic mtDNA polymorphism in the mitochondrial-derived peptide, MOTS-c. Aging (2021). 10.18632/aging.202529

124 Barman, P. K. et al. Production of <scp>MHCII</scp> -expressing classical monocytes increases during aging in mice and humans. Aging cell 21 (2022). 10.1111/acel.13701

125 Almanzar, N. et al. A single-cell transcriptomic atlas characterizes ageing tissues in the mouse. Nature 583, 590–595 (2020). 10.1038/s41586-020-2496-1

126 Teo, Y. V., Hinthorn, S. J., Webb, A. E. & Neretti, N. Single-cell transcriptomics of peripheral blood in the aging mouse. Aging 15, 6–20 (2023). 10.18632/aging.204471

127 Kimmel, J. C. et al. Murine single-cell RNA-seq reveals cell-identity- and tissue-specific trajectories of aging. Genome research 29, 2088–2103 (2019). 10.1101/gr.253880.119

128 Benayoun, B. A. et al. Remodeling of epigenome and transcriptome landscapes with aging in mice reveals widespread induction of inflammatory responses. Genome research 29, 697–709 (2019). 10.1101/gr.240093.118

129 Sun, Y. & Shang, D. Inhibitory Effects of Antimicrobial Peptides on Lipopolysaccharide-Induced Inflammation. 2015, 1–8 (2015). 10.1155/2015/167572

130 Tyers, M. & Wright, G. D. Drug combinations: a strategy to extend the life of antibiotics in the 21st century. Nature Reviews Microbiology 17, 141–155 (2019). 10.1038/s41579-018-0141-x

131 Ming, W. et al. Mitochondria related peptide MOTS-c suppresses ovariectomy-induced bone loss via AMPK activation. Biochemical and biophysical research communications 476, 412–419 (2016). 10.1016/j.bbrc.2016.05.135

132 Hu, B. T. & Chen, W. Z. MOTS-c improves osteoporosis by promoting osteogenic differentiation of bone marrow mesenchymal stem cells via TGF-beta/Smad pathway. Eur Rev Med Pharmacol Sci 22, 7156–7163 (2018). 10.26355/eurrev_201811_16247

133 Che, N. et al. MOTS-c improves osteoporosis by promoting the synthesis of type I collagen in osteoblasts via TGF-β/SMAD signaling pathway. European review for medical and pharmacological sciences 23, 3183–3189 (2019).

134 Kim, S. J., et al. The mitochondrial-derived peptide MOTS-c is a regulator of plasma metabolites and enhances insulin sensitivity. Physiological Reports 7 (2019). 10.14814/phy2.14171

135 Li, Q. et al. Earlier changes in mice after D-galactose treatment were improved by mitochondria derived small peptide MOTS-c. Biochemical and biophysical research communications (2019). 10.1016/j.bbrc.2019.03.194

136 Liu, C. et al. Reduced skeletal muscle expression of mitochondrial-derived peptides humanin and MOTS-C and Nrf2 in chronic kidney disease. American Journal of Physiology-Renal Physiology 317, F1122–F1131 (2019). 10.1152/ajprenal.00202.2019

137 Lu et al. Mitochondrial-Derived Peptide MOTS-c Increases Adipose Thermogenic Activation to Promote Cold Adaptation. International journal of molecular sciences 20, 2456 (2019). 10.3390/ijms20102456

138 Lu, H. et al. MOTS-c peptide regulates adipose homeostasis to prevent ovariectomy-induced metabolic dysfunction. J Mol Med (Berl) (2019). 10.1007/s00109-018-01738-w

139 Weng, F. B. et al. MOTS-c accelerates bone fracture healing by stimulating osteogenesis of bone marrow mesenchymal stem cells via positively regulating FOXF1 to activate the TGF-beta pathway. Eur Rev Med Pharmacol Sci 23, 10623–10630 (2019). 10.26355/eurrev_201912_19759

140 Yan, Z. et al. MOTS-c inhibits Osteolysis in the Mouse Calvaria by affecting osteocyte-osteoclast crosstalk and inhibiting inflammation. Pharmacological Research 147, 104381 (2019). 10.1016/j.phrs.2019.104381

141 Wei, M. et al. Mitochondrial-Derived Peptide MOTS-c Attenuates Vascular Calcification and Secondary Myocardial Remodeling via Adenosine Monophosphate-Activated Protein Kinase Signaling Pathway. Cardiorenal Med 10, 42–50 (2020). 10.1159/000503224

142 Xinqiang, Y., Quan, C., Yuanyuan, J. & Hanmei, X. Protective effect of MOTS-c on acute lung injury induced by lipopolysaccharide in mice. International immunopharmacology 80, 106174 (2020). 10.1016/j.intimp.2019.106174

143 Yin, X. et al. The intraperitoneal administration of MOTS-c produces antinociceptive and anti-inflammatory effects through the activation of AMPK pathway in the mouse formalin test. European journal of pharmacology 870, 172909 (2020). 10.1016/j.ejphar.2020.172909

144 Le, C.-F., Fang, C.-M. & Sekaran, S. D. Intracellular Targeting Mechanisms by Antimicrobial Peptides. Antimicrobial Agents and Chemotherapy 61 (2017). 10.1128/aac.02340-16

145 Quirós, P. M., Mottis, A. & Auwerx, J. Mitonuclear communication in homeostasis and stress. Nature Reviews Molecular Cell Biology 17, 213–226 (2016). 10.1038/nrm.2016.23

146 Sloan, D. B. et al. Cytonuclear integration and co-evolution. Nature Reviews Genetics 19, 635–648 (2018). 10.1038/s41576-018-0035-9

147 Reynolds, J. C., Bwiza, C. P. & Lee, C. Mitonuclear genomics and aging. Human Genetics 139, 381–399 (2020). 10.1007/s00439-020-02119-5

148 Tan, J. X. & Finkel, T. Mitochondria as intracellular signaling platforms in health and disease. Journal of Cell Biology 219 (2020). 10.1083/jcb.202002179

149 Gubert, C. & Hannan, A. J. Exercise mimetics: harnessing the therapeutic effects of physical activity. Nature Reviews Drug Discovery (2021). 10.1038/s41573-021-00217-1

150 Zhai, D. et al. MOTS-c peptide increases survival and decreases bacterial load in mice infected with MRSA. Mol Immunol 92, 151–160 (2017). 10.1016/j.molimm.2017.10.017

151 Gomes, B. et al. Designing improved active peptides for therapeutic approaches against infectious diseases. Biotechnology advances 36, 415–429 (2018). 10.1016/j.biotechadv.2018.01.004

152 Mishra, B., Reiling, S., Zarena, D. & Wang, G. Host defense antimicrobial peptides as antibiotics: design and application strategies. Current opinion in chemical biology 38, 87–96 (2017). 10.1016/j.cbpa.2017.03.014

153 Gagnon, K. T., Li, L., Janowski, B. A. & Corey, D. R. Analysis of nuclear RNA interference in human cells by subcellular fractionation and Argonaute loading. Nature protocols 9, 2045–2060 (2014). 10.1038/nprot.2014.135

154 Bray, N. L., Pimentel, H., Melsted, P. & Pachter, L. Near-optimal probabilistic RNA-seq quantification. Nature biotechnology 34, 525–527 (2016). 10.1038/nbt.3519

155 Love, M. I., Huber, W. & Anders, S. Moderated estimation of fold change and dispersion for RNA-seq data with DESeq2. Genome biology 15 (2014). 10.1186/s13059-014-0550-8

156 Kolde, R. R: pheatmap. 2015-12-11. R Package version 1.0.10. . (2015).

157 Subramanian, A. et al. Gene set enrichment analysis: a knowledge-based approach for interpreting genome-wide expression profiles. Proceedings of the National Academy of Sciences of the United States of America 102, 15545–15550 (2005). 10.1073/pnas.0506580102

158 Lu, R., Sampathkumar, N. K. & Benayoun, B. A. Measuring Phagocytosis in Bone Marrow-Derived Macrophages and Peritoneal Macrophages with Aging. Methods in molecular biology 2144, 161–170 (2020). 10.1007/978-1-0716-0592-9_14

159 Amend, S. R., Valkenburg, K. C. & Pienta, K. J. Murine Hind Limb Long Bone Dissection and Bone Marrow Isolation. J Vis Exp (2016). 10.3791/53936

160 Farouni, R., Djambazian, H., Ferri, L. E., Ragoussis, J. & Najafabadi, H. S. Model-based analysis of sample index hopping reveals its widespread artifacts in multiplexed single-cell RNA-sequencing. Nature communications 11, 2704 (2020). 10.1038/s41467-020-16522-z

161 Stuart, T. et al. Comprehensive Integration of Single-Cell Data. Cell 177, 1888–1902.e1821 (2019). 10.1016/j.cell.2019.05.031

162 McGinnis, C. S., Murrow, L. M. & Gartner, Z. J. DoubletFinder: Doublet Detection in Single-Cell RNA Sequencing Data Using Artificial Nearest Neighbors. Cell Syst 8, 329–337.e324 (2019). 10.1016/j.cels.2019.03.003

