## Supplemental Figures for "The Human Mitochondrial Genome Encodes for an Interferon-Responsive Host Defense Peptide"

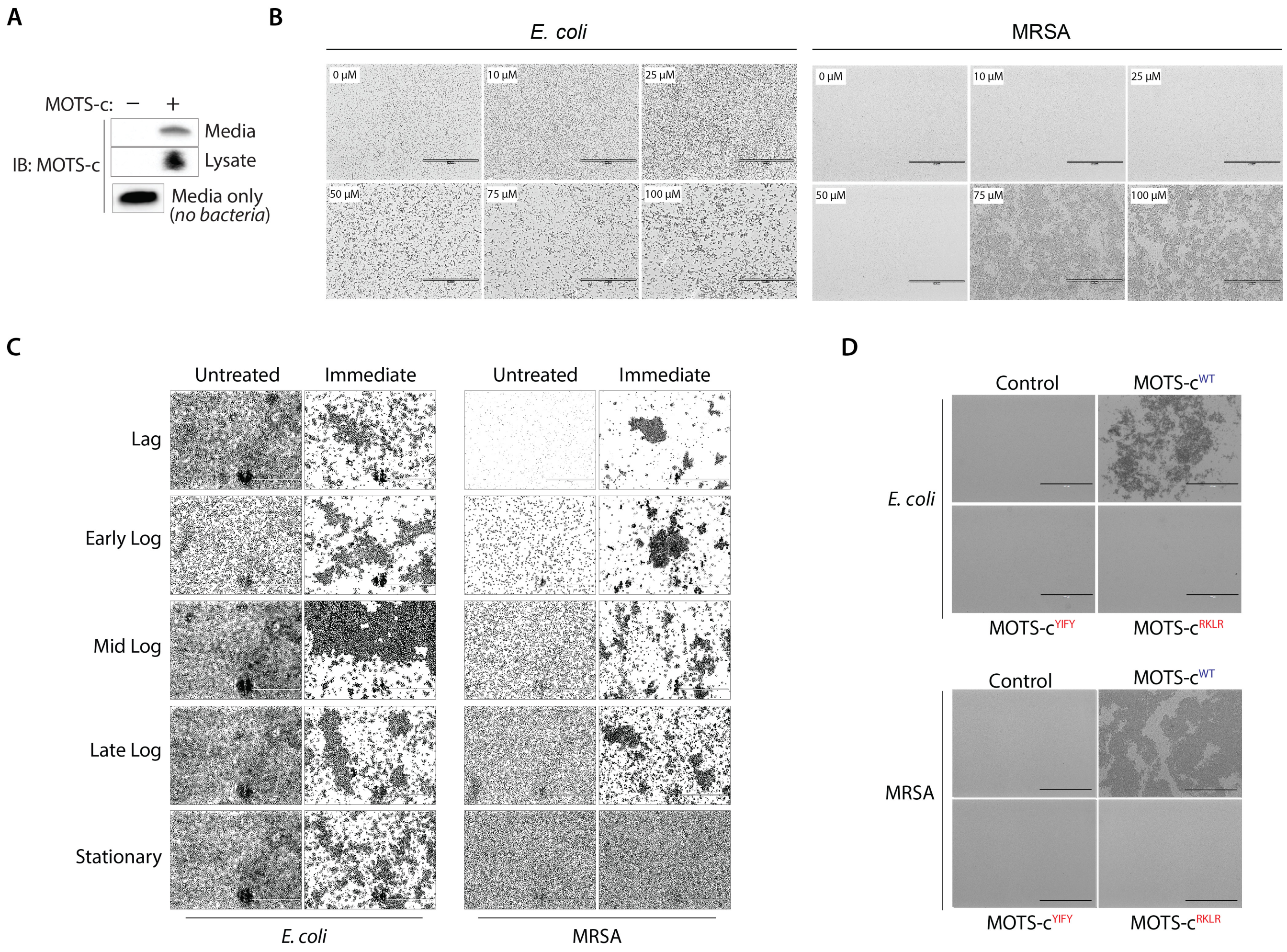


**Figure S1. The effect of MOTS-c bacterial aggregation.**

(**A**) MOTS-c (100 μM) was added to an *E. coli* suspension in media (water), which immediately aggregated and precipitated bacteria. MOTS-c levels remaining in media (water) and associated with *E. coli* were detected by Western blotting. MOTS-c (100 μM) levels in media (water) without any bacteria is also shown as reference. (**B**) MOTS-c immediately agglutinates bacteria in a dose-dependent manner. *E. coli* and MRSA were resuspended in water and treated with MOTS-c at varying doses (n=6). Bar, 200 μm. (**C**) *E. coli* and MRSA were collected at various points of growth (i.e. lag, early/mid/late log, and stationary phase), washed and resuspended in water, then treated with MOTS-c (100 μM). Images were taken immediately and rendered equally across all panels to aid in visual recognition of bacteria (n=3). Bar, 200 μm. (**D**) *E. coli* and MRSA were treated with WT or mutant MOTS-c (100 μM) and imaged immediately (n=6). MOTS-c mutants were devoid of its hydrophobic ($_{8}$YIFY$_{11}$>$_{8}$AAAA$_{11}$; MOTS-c$^{YIFY}$) or cationic domain ($_{13}$RKLR$_{16}$>$_{13}$AAAA$_{16}$; MOTS-c$^{RKLR}$).

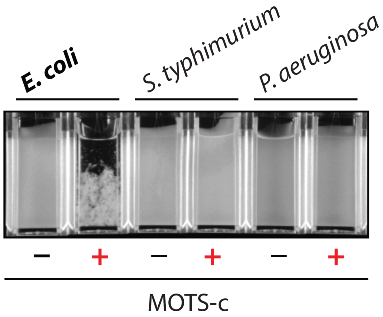


**Figure S2. MOTS-c exhibits specificity in bacterial targeting.** MOTS-c (100 μM) treatment causes immediate aggregation of *E. coli* but not *S. typhimurium* or *P. aeruginosa* (n=6). Representative image shown. See Figure 1D.

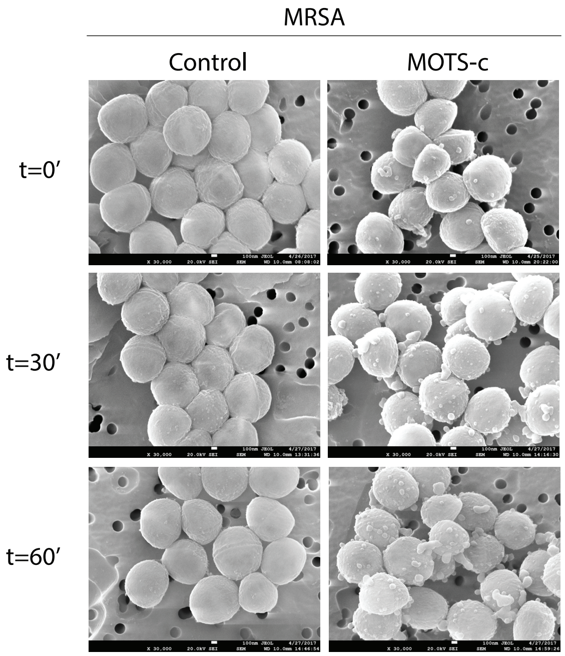


**Figure S3. MOTS-c targets MRSA membranes**. Scanning electron micrographs of methicillin-resistant *S. aureus* (MRSA) treated with MOTS-c (100 μM) for 0 (immediate fixation), 30, and 60 minutes (n=3). Representative images shown. Bar, 100 nm.

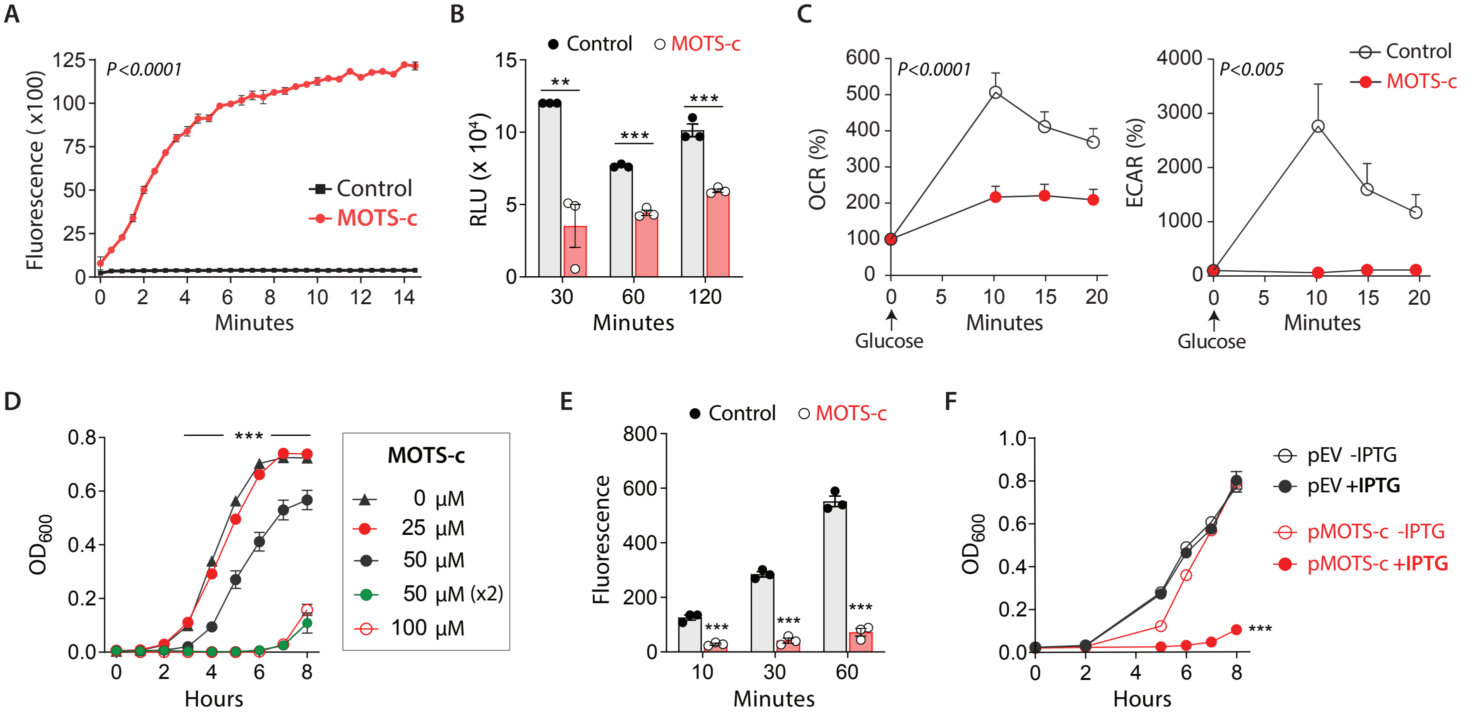


**Figure S4. The effect of MOTS-c on bacterial metabolism and growth.**
(**A**) Membrane integrity assessment by time-course quantification of SYTOX Green nucleic acid stain, which is excluded by intact *E. coli* membranes (n=6). (**B**) Total cellular ATP levels measured following MOTS-c treatment (100 μM) in *E. coli* (n=3). RLU: relative light units. (**C**) Real-time metabolic assessment of *E. coli* respiration (oxygen consumption rate; OCR) and glycolysis (extracellular acidification rate; ECAR)(n=8). (**D**) Growth curve of *E. coli*, measured by optical density at 600 nm ($OD_{600}$) following various doses of MOTS-c peptide treatment (n=6), (**E**) *E. coli* growth in M9 media with/without MOTS-c treatment (100 μM) was assessed using alamarBlue, a redox-sensitive resazurin dye (n=3). RFU: relative fluorescence units. (**F**) *E. coli* were transformed with expression vectors harboring the MOTS-c ORF (open reading frame) that can be induced by IPTG (Isopropyl β-D-1-thiogalactopyranoside). Growth determined by optical density at 600 nm ($OD_{600}$) periodically for 8 hours (n=3). Data expressed as mean +/- SEM. Mann-Whitney test for (B, E) and Two-way ANOVA (repeated measures) (A, C, D, and F). ** P<0.01 *** P<0.001.

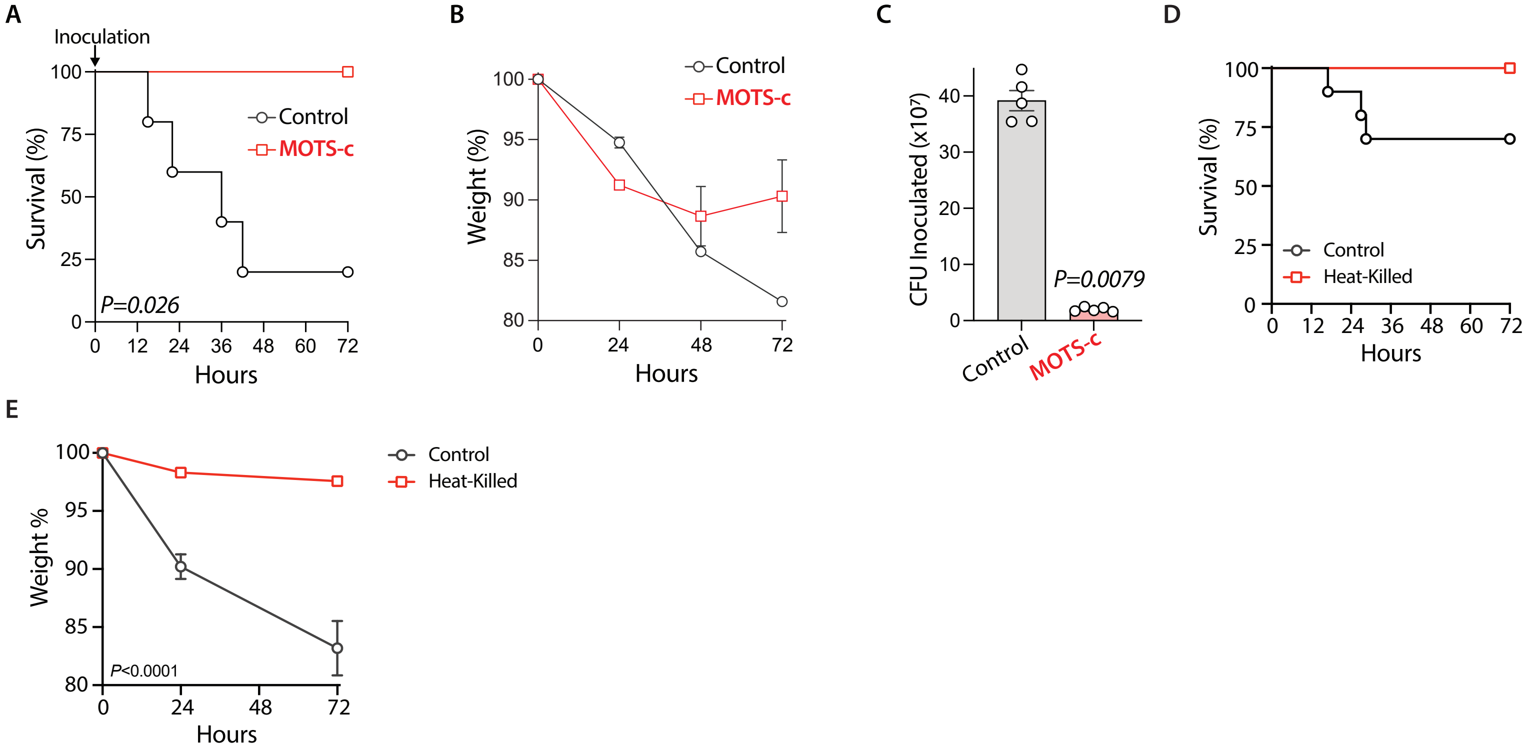


**Figure S5. MOTS-c enhances survival from MRSA exposure *in vivo* in male mice.**
(**A-C**) 4x$10^8$ CFU of mid-log phase MRSA was resuspended in 100 µM MOTS-c or vehicle (water) and immediately injected IP into 3.5-4 month-old male C57BL/6J mice (n=4-5). Survival (**A**) and weight (**B**) were monitored for 72 hours. (**C**) 4x$10^8$ CFU of MRSA resuspended in 100 µM MOTS-c or water was serial diluted and plated on LB agar before injection and colonies counted after overnight incubation. (**D-E**) 4x$10^8$ CFU of mid-log phase *S. aureus* (MRSA) was resuspended in phosphate-buffered saline (PBS) and either kept on ice (control) or heat-killed in a 70˚C water bath for 20 minutes. Live or heat-killed MRSA was injected IP into 3.5-4 month-old male C57BL/6J mice (n=10). Survival (**D**) and weight (**E**) were monitored for 72 hours. Data expressed as mean +/- SEM. Log-rank (Mantel-Cox) test (A,D), Mann-Whitney test (C), and 2-way ANOVA (repeated measures)(E).

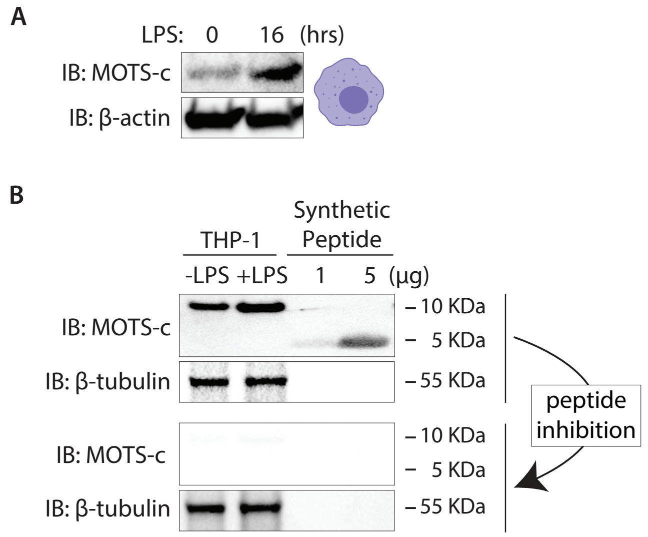


**Figure S6. Endogenous MOTS-c levels in macrophages.**
(**A**) Total endogenous MOTS-c levels in differentiated macrophages (THP-1) measured following LPS treatment for 16 hours. (**B**) Comparison of endogenous MOTS-c levels in macrophages (THP-1; 15 μg total cell lysate/well) and measured synthetic peptide levels (1 and 5 μg). Antibodies were competed out with MOTS-c peptide to confirm detection specificity. Synthetic MOTS-c peptide was chemically synthesized without any post-translational modifications (PTM) or oligomerizations.

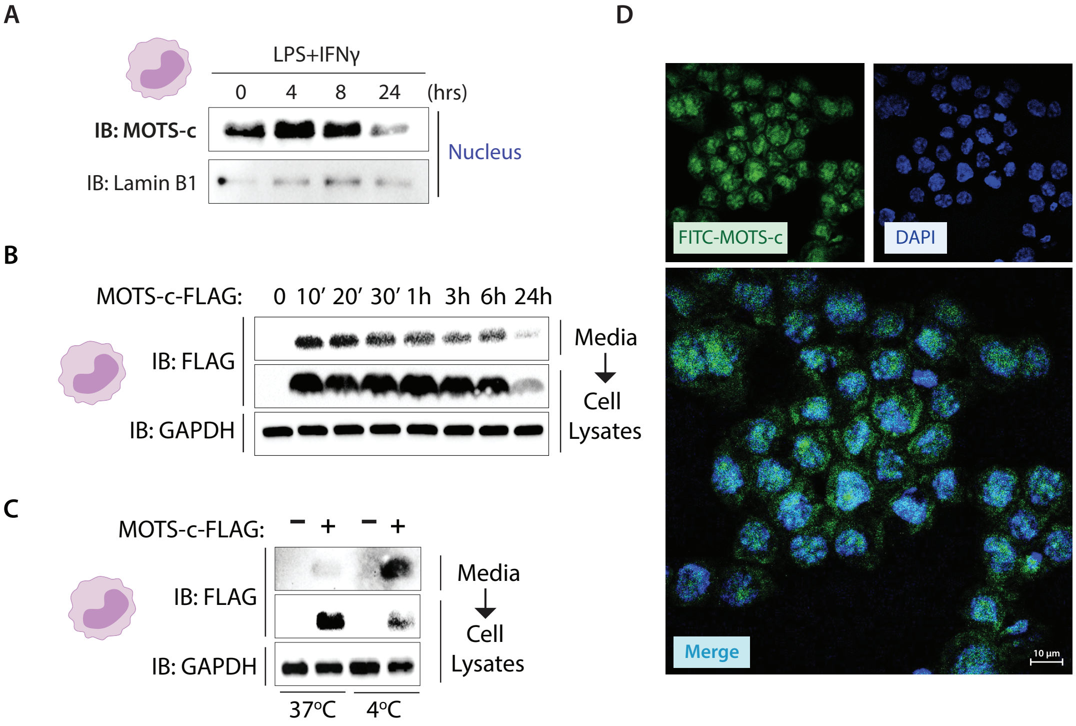


**Figure S7. Nuclear MOTS-c in monocytes during activation and exogenous MOTS-c treatment.** (**A**) MOTS-c translocates to the nucleus upon activation by LPS+IFNγ. A time-course measurement of endogenous MOTS-c in purified nuclear extracts following monocyte (THP-1) activation by LPS+IFNγ. Lamin B1 was used as a loading control for purified nuclear samples. See also Figure 4A. (**B**) Time-course detection of intracellular MOTS-c-FLAG peptide (10 μM) following treatment in THP-1 monocytes. (**C**) Intracellular levels of MOTS-c-FLAG peptide (10 μM) following treatment in THP-1 monocytes at different temperatures (i.e. 4˚C vs. 37˚C) for 30 minutes. (**D**) Exogenously treated MOTS-c peptide enters monocytes and localizes to the nucleus. Confocal fluorescence images of monocytes (THP-1) treated with FITC-MOTS-c (1 μM) for 30 minutes, showing nuclear localization. Nucleus marked by DAPI staining. Bar, 10 μm.

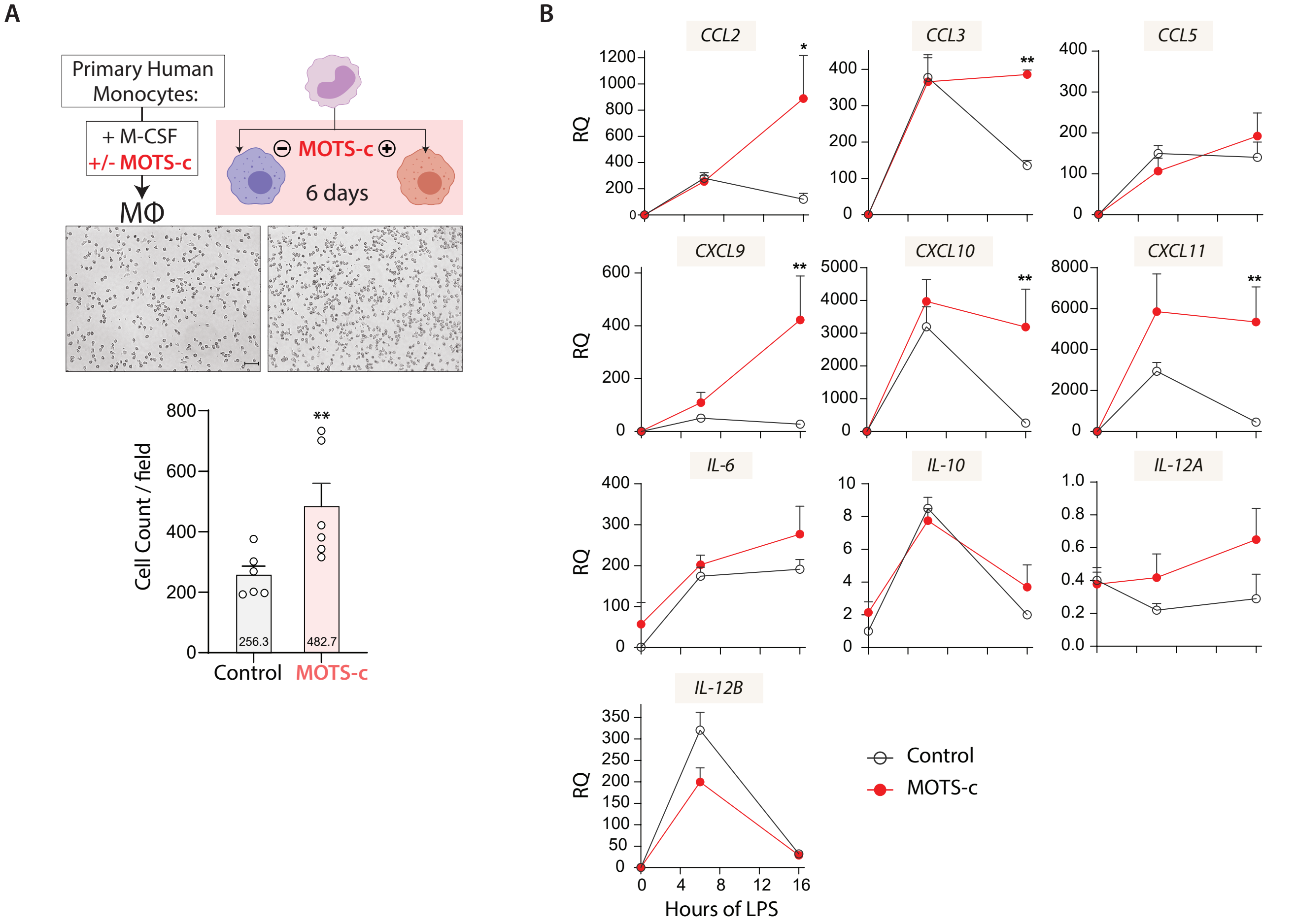


**Figure S8. The presence of MOTS-c during monocyte-derived macrophage differentiation generates macrophages with an altered phenotype.**

(**A**) MOTS-c affects primary human monocyte differentiation. Primary human monocytes were differentiated with M-CSF for 6 days with MOTS-c (10 μM) treatment or vehicle (dd$H_2O$) only during the first 24 hours (n=6; MΦ=macrophage; Bar, 10 μm). Adherent macrophages were imaged and counted. See also Figure 5A. (**B**) Macrophages (THP-1) were differentiated with MOTS-c (10 μM) or vehicle (dd$H_2O$) (n=6) for 4 days. Time-course gene expression levels in MOTS-c-programmed macrophages by qPCR of cytokines *CCL2, CCL3, CCL5, CXCL9, CXCL10, CXCL11, IL-6, IL-10, IL-12A*, and *IL-12B* following LPS treatment (100ng/ml)(0, 6, and 16 hours). Relative gene expression, calculated based on the 0 timepoint, is shown. RQ: relative quantification. See also Figure 5D. Data expressed as mean +/- SEM. Mann-Whitney test, * $P<0.05$, ** $P<0.01$.

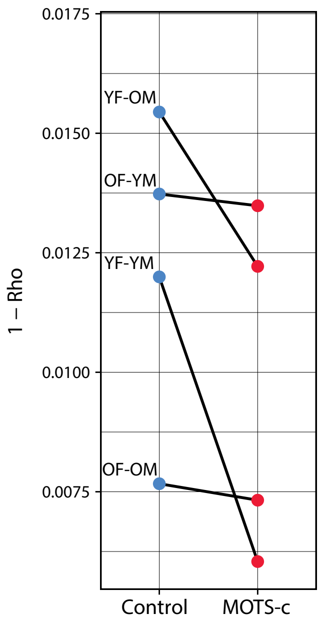


**Figure S9. Pairwise distance analysis between female and male samples in control and MOTS-c treated conditions for BMDM pseudobulk transcriptomes.**
The distance between each pair of (Female, Male) pseudobulk sample is computed as 1-Rho for each pair (YF-YM, YF-OM, OF-YM and OF-OM) in the control and MOTS-c treated conditions. To test whether the distance between (Female, Male) pseudobulk sample pair is reduced upon MOTS-c treatment, we used a one-sided paired Wilcoxon rank sum test ($p \sim 0.0625$). YF=young female; YM=young male; OF=old female; OM=old male.

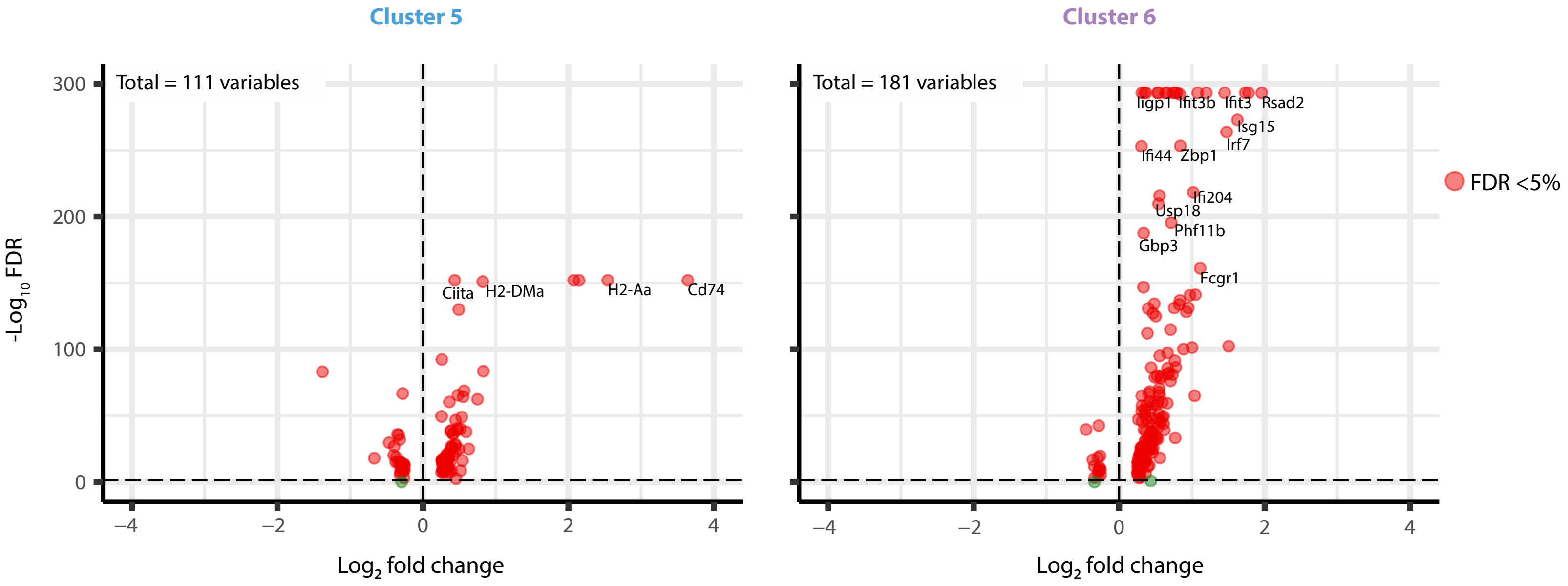


**Figure S10. Genes enriched in MOTS-c-programmed macrophage populations.**

Single-cell RNA-seq (scRNA-seq) was performed on bone marrow-derived macrophages (BMDMs) from young (4 mo.) and old (20 mo.) mice of both sexes that were differentiated for 7 days with MOTS-c (10 μM) or vehicle (dd$H_2O$). MOTS-c was given only once concomitantly with M-CSF at the onset of differentiation, and the media was replaced after 3 days in both the control- and MOTS-c-treated conditions (see also Figure 6A). Volcano plots on differentially expressed genes in clusters 5 and 6 that are enriched by MOTS-c are shown here (see also Figures 6D-6F, S11-S12 and Table S3).

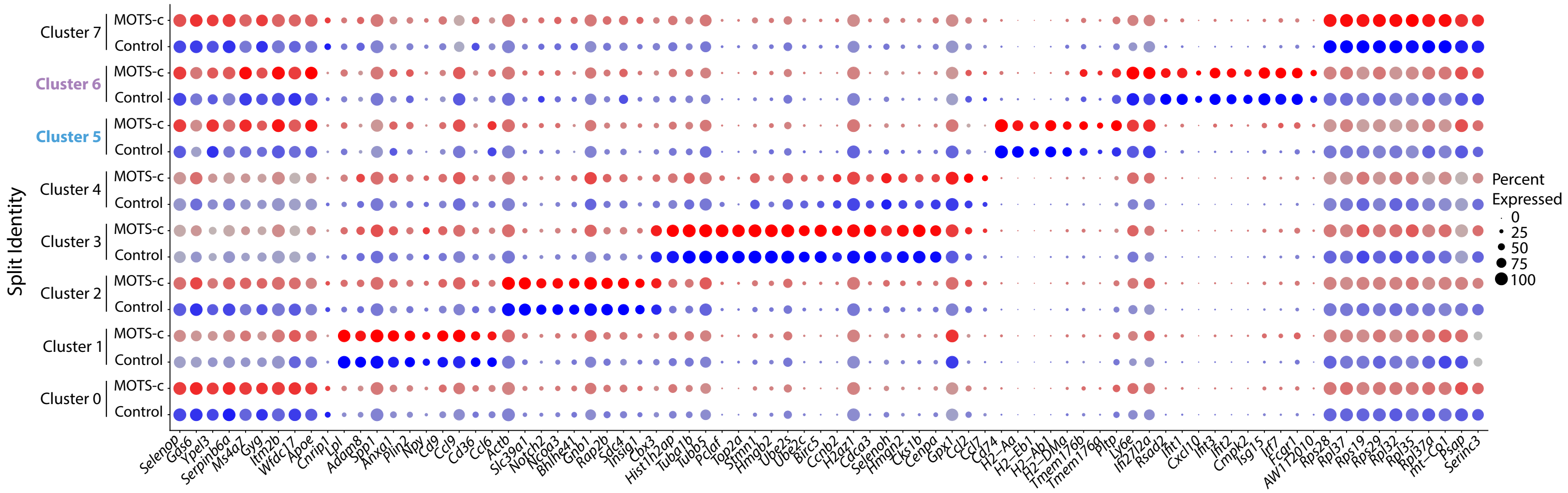


**Figure S11. MOTS-c generates macrophages that uniquely express genes related to antigen presentation and interferon signaling.** Single-cell RNA-seq (scRNA-seq) was performed on bone marrow-derived macrophages (BMDMs) from young (4 mo.) and old (20 mo.) mice of both sexes that were differentiated for 7 days with MOTS-c (10 uM) or vehicle (ddH$_2$O). MOTS-c was given only once concomitantly with M-CSF at the onset of differentiation, and the media was replaced after 3 days in both the control- and MOTS-c-treated conditions (see also Figure 6A). Dotplot of genes enriched in each of the 8 clusters, showing MOTS-c-programmed BMDMs (clusters 5 & 6) have a unique gene expression profile not present in other clusters (see also Figures 6D-6F, S10, S12 and Table S3).

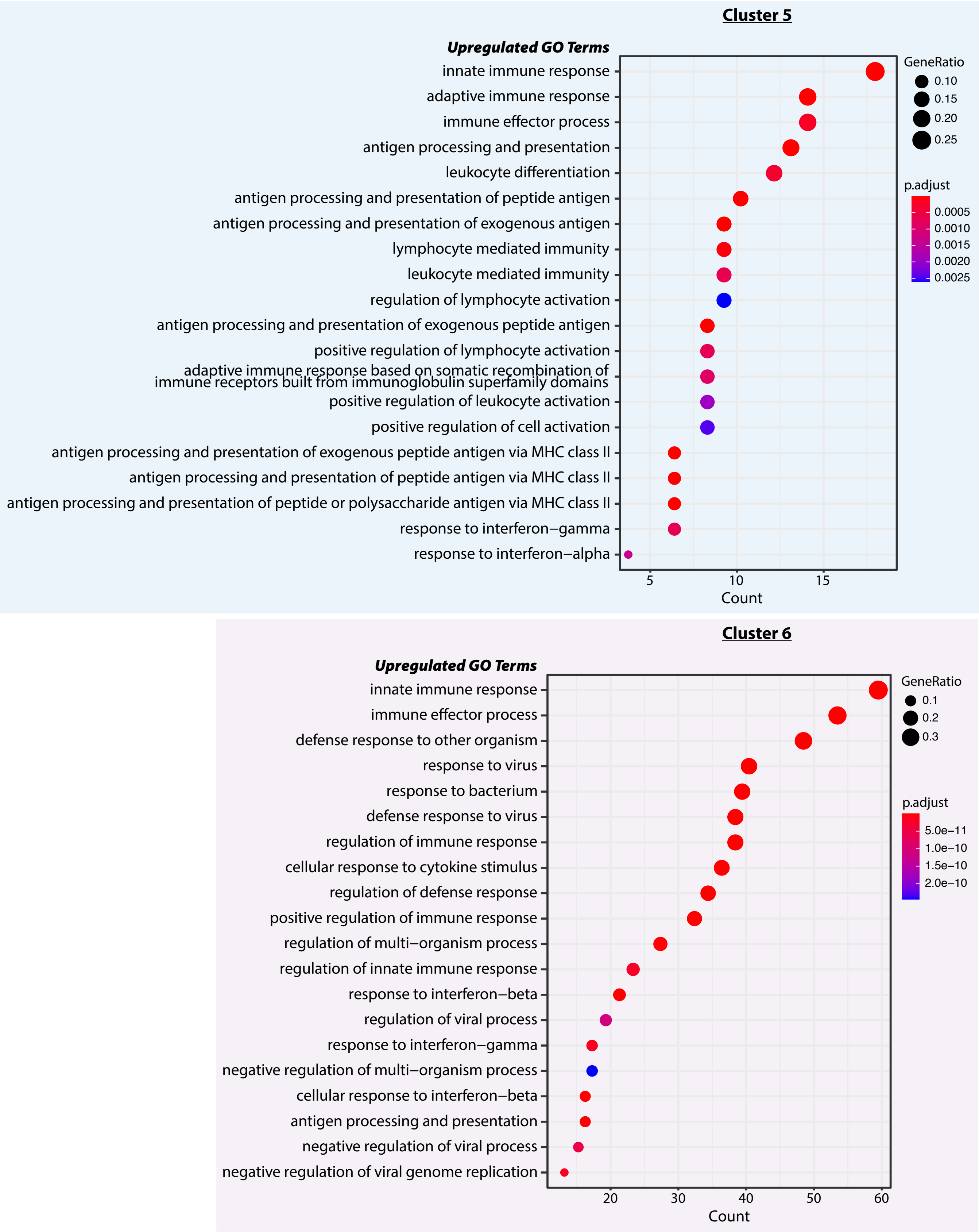


**Figure S12. MOTS-c-programmed macrophage populations exhibit differential regulation of antigen presentation and interferon-related pathways.**

Single-cell RNA-seq (scRNA-seq) was performed on bone marrow-derived macrophages (BMDMs) from young (4 mo.) and old (20 mo.) mice of both sexes that were differentiated for 7 days in the presence/absence of MOTS-c (10 µM). MOTS-c was given only once concomitantly with M-CSF at the onset of differentiation, and the media was replaced after 3 days in both the control- and MOTS-c-treated conditions (see also Figure 6A). Dotplots of biological processes derived from scRNA-seq data for clusters 5 and 6, based on gene set enrichment analysis (GSEA) of Gene Ontology Biological Process (GO_BP) at FDR < 5%. (see also Figures 6D-6F, S10-S11 and Table S3).
